# ATF4 Coordinates Transcriptomic and Structural Adaptations in Aging Muscle

**DOI:** 10.64898/2026.03.27.711928

**Authors:** Amber Crabtree, Mohd Mabood Khan, Estevão Scudese, Calixto Pablo Hernandez Perez, Prasanna Venkhatesh, Andrea G. Marshall, Benjamin Rodriguez, Edgar Garza López, Okwute M. Ochayi, Estélio Henrique Martin Dantas, Pamela Martin, Matheus Baffi, Fabiana Scartoni, Margaret Mungai, Kit Neikirk, Jennifer Streeter, Renata O. Pereira, Dao Fu Dai, Han Le, Harrison Mobley, Jeremiah Afolabi, Bret C. Mobley, Celestine N. Wanjalla, Duane Hall, Julia D. Berry, Oleg Kovtun, Jenny C. Schafer, Sean Schaffer, Prasanna Katti, Chantell Evans, André Kinder, Joyonna Gamble-George, Melanie McReynolds, Annet Kirabo, Sepiso K. Masenga, Antentor O. Hinton

## Abstract

Aging is associated with a progressive loss of skeletal muscle function, known as sarcopenia; however, the molecular mechanisms coordinating cellular stress responses and structural adaptations permissive of sarcopenia remain incompletely understood. In our previous studies, we found aging differentially impacted mitochondrial networks by muscle, suggesting unique stress thresholds and response activation. Here, we investigate the role of activating transcription factor 4 (ATF4), a master regulator of the integrated stress response (ISR), in aged quadriceps muscle using complementary patient and aging mouse models. Older adults exhibited a marked decrease in aerobic capacity, muscle strength, and endurance when compared with young participants. These results paralleled findings in aged mice, with significant loss of muscle mass across multiple hindlimb muscles. Ultrastructural analysis revealed substantial age-related changes in mitochondrial morphology, including increased volume, surface area, and branching index, as well as a shift toward larger, more complex mitochondria. Our data indicate that ATF4 binds directly to the promoter region of the gene encoding TFAM, suggesting a transcriptional regulatory relationship to support DNA stability. These structural and transcriptional changes likely impair oxidative capacity and drive a feed-forward cycle of mitochondrial dysfunction and ISR activation. Our findings indicate that ATF4 coordinates transcriptomic and structural adaptations in aging muscle, identifying the ISR pathway as a potential therapeutic target for preserving muscle function in older adults.

## Introduction

Aging is accompanied by a progressive loss of skeletal muscle mass, strength, and functional capacity known as sarcopenia. Sarcopenia impairs quality of life and increases susceptibility to metabolic and cardiovascular diseases, making it a major contributor to age-related morbidity (Adams, Ebert et al. 2017, Mao, Lv et al. 2025). Although sarcopenia is associated with mitochondrial dysfunction, metabolic stress, hormonal fluctuations, and mechanical alterations (Lang, Streeper et al. 2010), the full molecular pathways that integrate these stresses to induce maladaptive muscle remodeling with age are not fully known. Two key contributors to mitochondrial decline under stress are disruptions in proteostasis, the system responsible for maintaining proper protein folding and turnover, and damage to mitochondrial DNA (mtDNA) and structure, which impairs energy producing processes within the cell (Vasudevan, Neuman et al. 2020).

As organisms age, the accumulation of misfolded proteins and mtDNA mutations increases cellular stress and activates the integrated stress response (ISR), a protective signaling pathway that attempts to restore cellular homeostasis. The ISR is initiated by the phosphorylation of the eukaryotic initiation factor 2 alpha subunit (eIF2α) by one of four kinases that respond to stress: General Control Nonderepressible 2 (GCN2), Protein Kinase R-like ER Kinase (PERK), Heme Regulated Inhibitor (HRI), or Protein Kinase R (PKR). This process leads to a decrease in global protein synthesis, providing a means for the cell to overcome the burden of the imposed stress, while simultaneously promoting the translation of activating transcription factor 4 (ATF4) (Vasudevan, Neuman et al. 2020), which is recognized as the main ISR effector. This selective translation is regulated by upstream open reading frames in the 5′ untranslated region of ATF4 mRNA and is further supported by atypical initiation factors, such as eIF2D and DENR, which allow ATF4 synthesis to occur even when translational repression is sustained (Ait Ghezala, Jolles et al. 2012).

ATF4 serves as a central integrator of cellular stress responses, linking proteostasis to mitochondrial function and genome maintenance (Memme, Mesbah Moosavi et al. 2020). Under conditions of proteotoxic or metabolic stress, activation of ATF4 through the integrated stress response promotes the expression of genes involved in protein folding, quality control, and degradation pathways, thereby restoring proteostasis. Importantly, this adaptive program extends to the mitochondria, where proper proteostasis is essential for maintaining mitochondrial protein import, electron transport chain integrity, and overall organelle function (Quirós, Prado et al. 2017). ATF4-dependent signaling has also been implicated in regulating mitochondrial transcriptional networks that support mtDNA stability, replication, and distribution. Disruption of proteostasis can lead to accumulation of misfolded mitochondrial proteins, impaired respiratory capacity, and mtDNA instability, whereas ATF4 activation helps coordinate a compensatory response that preserves mitochondrial integrity (Harding, Zhang et al. 2003, Walter and Ron 2011). Thus, ATF4 functions as a key node that couples protein homeostasis with mitochondrial maintenance, ensuring proper bioenergetics and cellular adaptation during stress.

ATF4 functions as a master regulator of amino acid metabolism, one-carbon metabolism, redox balance, and stress adaptation (Kreß, Jessen et al. 2023). In skeletal muscle, ATF4 signaling plays a dual and context-dependent role. Experimental studies demonstrate that ATF4 is both sufficient and required to induce muscle fiber atrophy during aging, fasting, and disuse, in part through transcriptional activation of atrophy-associated genes such as *Gadd45a* and *Cdkn1a* (Adams, Ebert et al. 2017, Miller, Marcotte et al. 2023). However, in the situations of acute stress, ATF4-mediated metabolic reprogramming can perform compensatory functions by preserving the availability of amino acids and antioxidant capability (Ebert, Dyle et al. 2015, Adams, Ebert et al. 2017).

Emerging evidence suggests that dysregulated or prolonged ISR activation contributes to age-related muscle dysfunction (Brown, Scaramozza et al. 2025). Prolonged ATF4 signaling has been connected to oxidative stress, mitochondrial remodeling, and enhanced susceptibility to aging (Brown, Scaramozza et al. 2025). Furthermore, ATF4 activity is closely regulated by feedback mechanisms involving downstream transcription factors like CHOP, which restrict excessive ISR signaling and maladaptive transcriptional outputs (Kaspar, Oertlin et al. 2021). Disruption of this regulatory balance may cause ATF4 signaling to change from adaptive to pathogenic with aging.

Research has shown changes in mitochondrial networks and proteins associated with mitochondrial fusion and cristae organization, including mitofusin-2 (MFN2) and components of the MICOS complex. These proteins contribute to the mitochondrial membrane structure and the appropriate organization of the mitochondrial network. Muscle function may be diminished as a result of age-related alterations in pathways connected to MFN2 and MICOS, which have the ability to decrease oxidative phosphorylation and damage mitochondrial connectivity and cristae structure (Vue, Garza-Lopez et al. 2023, Scudese, Marshall et al. 2025). Modifications in these processes can result in mitochondria that are fragmented or structurally aberrant, making them less efficient in producing energy. Modifications to mitochondrial shape and dynamics can impair energy metabolism and exacerbate oxidative damage, which in turn can affect cellular health and disease conditions (Hinton, Claypool et al. 2024, Hinton and Marshall 2024, Jenkins, Neikirk et al. 2024).

Importantly, ATF4 serves as a stress-responsive transcriptional regulator that integrates these age-related mitochondrial alterations by modulating genes involved in mitochondrial dynamics, proteostasis, and cristae organization, thereby linking cellular stress signaling to the maintenance or remodeling of mitochondrial structure during aging. In this work, we examine how ATF4 disruption affects gene regulatory networks, proteostasis, and mitochondrial architecture. We show that a crucial connection between cellular stress and mitochondrial genome stability is made via stress-responsive pathways, specifically ATF4 signaling. By combining transcriptome analysis, targeted suppression of the NGLY1– Nrf1 axis, and advanced imaging, we uncover a coordinated regulatory network that drives mitochondrial remodeling through abnormalities in proteostasis. Interestingly, in a mouse aging model, we found that ATF4 expression and mitochondrial morphology differ in the quadriceps compared to the gastrocnemius, indicating a fiber-type-specific response.

## Materials and Methods

### Study Population and Enrollment

The cohort had different specific recruitment criteria. According to predetermined criteria, older adults were chosen, which included being at least 50 years old, physically capable of exercising, and free of chronic disorders that could affect their ability to exercise or their muscle physiology.

This study’s “young” population was represented by a group of physical education students. The students’ ages varied from 18 to 50 years, and they were all registered in sports science and physical education courses as part of their educational requirements. All participants provided informed consent, and the study’s results were unaffected by any preexisting diseases or injuries. The experimental protocols followed internationally recognized human research norms, including the principles articulated by the World Medical Association in the Declaration of Helsinki, which establishes ethical standards for medical research involving human subjects (2013).

Exclusion criteria that were applied to cohort comprised the following: having a significant malignancy (such as solid tumors, hematological malignancies, or metastatic cancer), known co-morbidities that are not sarcopenia, being pregnant or planning to become pregnant during the study, having a significant cognitive impairment or psychiatric disorder that affects the capacity to give informed consent, having severe musculoskeletal injuries that limit mobility or exercise capacity, actively abusing substances, having recently participated in an intense physical training program, having surgery, and currently using medications that are known to significantly influence muscle anatomy or functioning, such as statins, neuromuscular blockers, and corticosteroids. The cohort characteristics are given Appendices S4.

This study used a variety of human cohorts from several different nations. All 3D reconstruction specimens were obtained from Vanderbilt University Medical Center. The Vanderbilt University Institutional Review Board (IRB) approved the collection of human tissue under the title “Mitochondria in Aging and Disease—Study of Archived and Autopsy Tissue” with the IRB number 231584. All other human samples were obtained from Brazilian cohorts using the CAEE (Ethics Appreciation Presentation Certificate) criteria. CAEE number 61743916.9.0000.5281 was used to collect samples from young people and conduct tests; CAEE number 10429819.6.0000.5285 was used to gather samples from older people. All studies included a combination of male and female samples, and the average age used as a cut-off for humans was around 50 years.

### Body Measurement and Physical Assessment

Body measurements, including height (cm), weight (kg), and body mass index (BMI; kg/m²), were recorded for all participants. Physical performance was assessed through several standardized tests. A 6-minute walk test was conducted on a level walking course, with participants instructed to walk as much as possible within the time limit. The total distance covered (meters) was recorded, and maximum oxygen consumption (VO₂ max) was estimated according to distance, age, body weight, and sex. Muscle strength was assessed bilaterally using a dynamometer to measure grip strength. The participant performed three maximal voluntary contractions for each hand, with 60-second rest intervals between trials. The highest value for each side was noted. Upper-body muscle endurance was evaluated using a chair stand test, with the number of complete sit-to-stand transitions recorded over 30 seconds.

### Animal Models and Tissue Collection

Male C57BL/6J mice at three different ages, 3 months, 1 year, and 2 years were used as models for young adult, middle-aged, and aged muscle, respectively (n = 5-7 per age group). We created Atf4^fl/fl^ mice following the methods mentioned earlier (Streeter, Persaud et al. 2024, Scudese, Marshall et al. 2025). A gracious donation from Dr. Pierre Chambon (University of Strasbourg) provided the tamoxifen-inducible HSA-CreERT2 mice. Atf4^fl/fl^ mice were crossed with HSA-CreERT2 mice to produce skeletal muscle-specific Atf4 knockout mice. The mice were housed under standard conditions with a 12-hour light-dark cycle and were provided with food and water ad libitum.

Mice that were 6 weeks old were given tamoxifen (20 mg/kg for males) intraperitoneally for 5 consecutive days to promote skeletal muscle recombination (Sigma, St. Louis, MO, USA; T5648). The mice were fed regular chow until four weeks following injection, and then the experiments were carried out according to the protocol. Mice were considered young adults for these investigations when they were 3–4 months in age, and old when they were 24–25 months in age when their tissues were harvested. There were no dietary, housing, or light-dark cycle changes between the age groups, and the aged animals were kept in the same environmental settings as the young cohorts and were checked for health issues.

The animals were euthanized via cervical dislocation followed by CO_2_ anesthesia. The time frame for collecting all serum and skeletal muscle samples was from 8:00 am to 12:00 pm. Additionally, body weight was recorded at the time of tissue harvest. Next, the hindlimb muscles, including quadriceps, plantaris, soleus, extensor digitorum longus (EDL), and gastrocnemius, were rapidly dissected and weighed to assess age-related changes in muscle mass. All procedures involving animal models were carried out in accordance with institutional animal care and use committee guidelines.

### GEO Dataset Analysis of ATF4 ChIP-seq

Analysis of publicly accessible GEO datasets (GSE35681) from Atf4^+/+^, ATF4^-/-^, Chop^+/+^, and Chop^-/-^ mouse embryonic fibroblasts (MEFs) following 10-hour tunicamycin (2 μg/mL) treatment that were studied using data deposited under GEO accession GSE35681, which was first described by Han et al (Han, Back et al. 2013) was performed. The datasets include ATF4 chromatin immunoprecipitation sequencing (ChIP-seq). We used the Galaxy platform to conduct all of our analyses. The ChIP-seq analysis began with Bowtie, which aligned the raw sequencing reads to the mouse reference genome. Using Model-based Analysis of ChIP-Seq (MACS), the Sequence Alignment Map (SAM) files were converted to the Binary Alignment Map (BAM) format, and peak calling was performed. The Integrated Genome Browser was used to view the identified ATF4 binding peaks.

### Serial block-face scanning electron microscopy (SBF-SEM)

Preparation of SBF-SEM was carried out as previously described (Vue, Garza-Lopez et al. 2023). Briefly, following a 5% isoflurane anesthesia, skin and hair were removed, the hindlimbs of male mice were immersed in a solution containing 2% glutaraldehyde and 100 mM phosphate buffer for a duration of 30 minutes. Gastrocnemius and quadriceps muscles were prepared by dissecting, chopping into 1 mm3 cubes, and soaking for 1 hour in a solution containing 2.5% glutaraldehyde, 1% paraformaldehyde, and 120 mM sodium cacodylate. After rinsing the tissues three times with 100 mM cacodylate buffer at room temperature, they were immersed in a solution containing 3% potassium ferrocyanide and 2% osmium tetroxide for 1 hour at 4°C. Samples were then treated with 0.1% thiocarbohydrazide and 2% filtered osmium tetroxide for 30 minutes. Finally, samples were kept overnight in a solution containing 1% uranyl acetate at 4°C.

Three washing with deionized water were carried out between each stage. After being submerged in a 0.6% lead aspartate solution for 30 minutes at 60°C the next day, the samples were dehydrated using acetone at varying concentrations. To dehydrate, we used acetone concentrations of 20%, 50%, 70%, 90%, 95%, and 100% for 5 minutes each. After being immersed in Epoxy Taab 812 strong resin, tissues were subjected to polymerization at 60°C for 36-48 hours. After polymerization, the blocks were sectioned for examination using a transmission electron microscope (TEM). The sections were then trimmed to a size of 0.5 mm × 0.5 mm and affixed to aluminum pins. Following previous methods (Vue, Garza-Lopez et al. 2023), the pins were run on an FEI/Thermo Scientific Volumescope 2 SEM, a cutting-edge SBF imaging machine, which produced 300–400 10 μm by 10 μm ultrathin (90nm) serial sections. Accordingly, with previous descriptions (Vue, Garza-Lopez et al. 2023), all sections were stained and photographed after being collected onto formvar-coated slot grids.

### Transmission Electron Microscopy

The samples used for ultrastructural studies were fixed in 2.5% glutaraldehyde in 0.1 M sodium cacodylate buffer at 4 °C and at a pH 7.4. Post-fixing was performed on fixed samples in 1% osmium tetroxide at room temperature for 1 hour. The dehydration process was performed using an alcohol series with concentrations of 30%, 50%, 70%, 95%, and 100%, each applied for one minute. Epoxy resin infiltrations at consecutive concentrations were performed, and the infiltrations were polymerized at 60°C for 48 hours.

Sections of Ultrathin (70-90nm) were cut using an ultramicrotome and collected on copper grids. The sections were stained for 10 minutes with 2% uranyl acetate, followed by 5 minutes with lead citrate to increase contrast. Transmission microscopy was performed at 5,000-20,000x magnification to observe mitochondrial ultrastructure along the muscle fibers (Lam, Katti et al. 2021, Neikirk, Lopez et al. 2023, Neikirk, Vue et al. 2023).

### Transmission Electron Microscopy (TEM) Analysis

TEM was carried out as previously described (Evers-van Gogh, Alex et al. 2015): Cells were fixed in 2.5% glutaraldehyde made in sodium cacodylate buffer for 1 hour at 37°C, then embedded in 2% agarose, post-fixed in 1% osmium tetroxide, dyed with 2% uranyl acetate, and dehydrated using a graded ethanol series. Following the embedding of samples in EMbed-812 resin, 80-nm slices were cut with an ultramicrotome and colored with uranyl acetate and lead citrate.

Images were obtained utilizing a JEOL JEM-1230 transmission electron microscope at 120 kV. Mitochondria and cristae were manually tracked and analyzed using the freehand tool in the ImageJ program (Parra, Moraga et al. 2013). Utilizing the Multi-Measure ROI tool (Vue, Garza-Lopez et al. 2023)The size, shape, and number of mitochondria were measured. Cristae volume was calculated as the ratio of total cristae area to total mitochondrial area, and cristae morphology was evaluated over three equivalent ROIs per image.

## ImageJ Quantification Workflow

- Quadrant selection using the ImageJ Quadrant Picking plugin
- Random selection of two quadrants per cell
- Minimum of 10 cells per individual across three individuals
- Expansion to 30 cells per individual if variability is observed
- Use of ROI Manager and specific measurement settings:

`o` Area
`o` Shape descriptors
`o` Perimeter
`o` Feret’s diameter
`o` Integrated density
`o` Fit ellipse

## Additional methods for TEM

Quantification analysis methods were developed using the National Institutes of Health (NIH) *ImageJ* software. *ImageJ* is an open-source image processing software designed to analyze multidimensional scientific images, such as TEM and confocal microscopy data sets. Notably, the NIH *ImageJ* software utilises pixel count to display an image in a 2048 x 2048-pixel frame. Each pixel in the frame is assigned a horizontal and vertical coordinate. Any straight line in the image can be defined by two pixels at its ends: pixel 1 and pixel 2.

Each pixel has Cartesian coordinates:

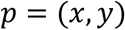

For two pixels:

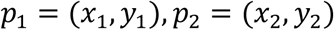

Measurements needed to decipher the changes in organelle morphology were constructed using the following calculations

**Length (*L*):** The length is measured by applying the Pythagorean Theorem to the coordinate pixels.

The straight-line distance between 𝑝_1_and 𝑝_2_is:

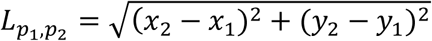

This is the standard Euclidean metric derived from the Pythagorean theorem.

**Area:** This is calculated similarly. The area calculated by ImageJ is the area of pixels.

**(a)** Rectangle

Let the side lengths be:

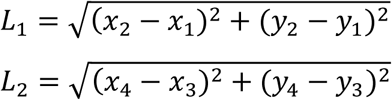

Then the rectangle area:

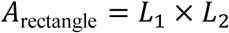

**(b)** Circles

Once the circle is traced, the software chooses two opposite pixels on the circle, pixel 1 and pixel 2

If two opposite pixels define the diameter 𝐿_𝐷_:

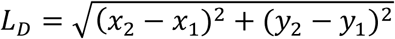

Radius:

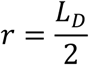

Circle area:

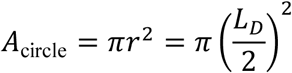

### Perimeter

Rectangle

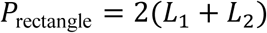

Circles

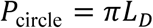

### Circularity Index (*Ci*)

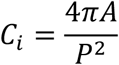

For a perfect circle:

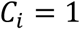

For irregular or elongated shapes:

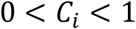

This metric is dimensionless because:

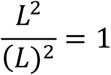

### Volume (2D-derived approximation)

If an estimate of volume is derived from surface-related measurements:

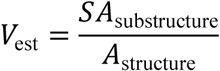

This is not a true geometric volume unless an axial depth term is incorporated. A physically meaningful volumetric estimate from 2D imaging typically requires multiplication by an assumed thickness 𝑡:

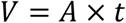

Otherwise, the ratio 𝑆𝐴: 𝐴 is a relative density metric, not a volume.

### ImageJ Parameters

First, a low-magnification image (in a DM3 or TIFF format) of the whole cell is uploaded to *ImageJ to quantify these many diverse measurements*. The entire cell is then divided into quadrants using the ImageJ plugin Quadrant Picking. https://imagej.nih.gov/ij/plugins/quadrant-picking/index.html

After splitting the first image into four quadrants, two quadrants must be randomly selected and used for each group’s complete analysis. To obtain accurate, reproducible values, these measurements should be repeated in at least 10 cells, with three analyses per cell. If there is variability across the three individuals’ data, per-individual values from 30 to 30 effectively reduce variability. During analysis, measurements can be tracked with the ROI Manager interface (Analyze > Tool > ROI Manager). Subsequently, the necessary measures can be set (Analyse> Set Measurements: Area, Mean grey value, Min & Max grey value, Shape descriptors, integrated density, Perimeter, Fit ellipse, Feret’s Diameter).

### Cristae Scoring and Morphometric Definitions

Cristae Score (0–4)

1. 0: No sharply defined cristae
2. 1: >50% of mitochondrial area without cristae
3. 2: >25% of mitochondrial area without cristae
4. 3: Many cristae (>75% of area) but irregular
5. 4: Many regular, well-organized cristae

**Cristae Surface Area**: Sum of the areas of all cristae within a single mitochondrion.

**Cristae Volume Density**: Cristae surface area divided by total mitochondrial area.

**Cristae Number**: Number of cristae per mitochondrion (either manually counted or derived from prior measurements).

3D parameters

### Quantification of Mitochondrial Volume, Surface Area, and Perimeter

3D mitochondrial reconstruction

Mitochondria were reconstructed from serial electron microscopy (EM) sections following manual or semi-automated segmentation of the outer mitochondrial membrane across consecutive slices. Segmented volumes were rendered as closed three-dimensional (3D) surface meshes in Amira (Thermo Fisher Scientific), enabling the extraction of quantitative morphometric parameters.

### Mitochondrial volume

Mitochondrial volume was calculated from the closed 3D surface mesh generated from serial EM reconstructions. Conceptually, volume corresponds to the total space enclosed by the triangulated surface and can be expressed as:

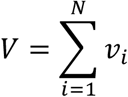

where

𝑣_𝑖_= volume of the 𝑖-th tetrahedral element

𝑁= total number of tetrahedral elements

Equivalently, when derived from voxel-based segmentation, mitochondrial volume can be represented as:

Mitochondrial Volume (voxel-based formulation)

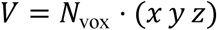

where

𝑁_vox_= number of voxels assigned to the mitochondrion

𝑥, 𝑦, 𝑧= calibrated voxel dimensions along each spatial axis Units: 𝑉in 𝜇𝑚^3^

### Mitochondrial surface area

Mitochondrial surface area was extracted directly from the 3D triangulated surface mesh generated following reconstruction. Surface area was calculated as the sum of the areas of all triangular faces composing the mesh

Mitochondrial Surface Area

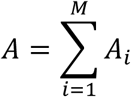

where

𝐴_𝑖_= area of the 𝑖-th triangular face

𝑀= total number of triangular faces

Units: 𝐴in 𝜇𝑚^2^

### Mitochondrial perimeter

Mitochondrial perimeter was quantified from two-dimensional (2D) EM cross-sections rather than from the 3D surface mesh. For each mitochondrion, the outer mitochondrial membrane was manually traced in representative EM sections, and the total contour length was calculated as

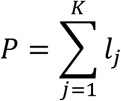

where

𝑙_𝑗_= length of the 𝑗-th contiguous segment

𝐾= total number of segments defining the contour

Units: 𝑃in 𝜇𝑚

### Derived shape metric (sphericity)

To assess mitochondrial shape independently of size, sphericity was calculated using mitochondrial volume and surface area according to the following equation:

The correct standard 3D sphericity equation is:

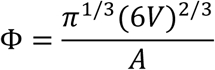

where

𝑉= mitochondrial volume

𝐴= mitochondrial surface area

Note carefully: surface area must be in the denominator. The earlier text inverted this. A perfect sphere yields Φ = 1.

### Mitochondrial Branching and Morphological Complexity (3DEM Parameters)

#### Conceptual definition

Mitochondrial branching and morphological complexity were quantified using a three-dimensional mitochondrial complexity index (MCI), a surface-to-volume-based metric designed to capture deviations from simple geometric shapes and reflect increased branching, elongation, and membrane convolutions arising from fusion events, nanotunnels, and network formation. Unlike two-dimensional form factor or aspect ratio measurements, MCI is derived from full three-dimensional reconstructions and is invariant to overall mitochondrial size, allowing direct comparison of structural complexity across conditions.

### Mitochondrial Complexity Index (MCI) Original dimensionless formulation

MCI was initially defined as a three-dimensional extension of the form factor metric, derived from surface area and volume measurements extracted from 3D mitochondrial reconstructions:

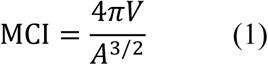

This is dimensionless because:

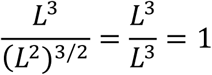

So, it is scale-invariant, as intended.

- A is mitochondrial surface area (μm²),
- V is mitochondrial volume (μm³).

The fractional exponent (3/2) ensures that the expression is dimensionless, as surface area scales with L^2^ and volume scales with L^3^. This formulation, therefore, removes dependence on absolute mitochondrial size and isolates morphological complexity. We confirmed that this formulation is volume-invariant and increases with mitochondrial branching and membrane irregularity.

### Expanded dynamic range formulation

Although Equation (1) reliably captures mitochondrial complexity, its scaling compresses the dynamic range and underrepresents perceived differences in highly branched or elongated mitochondria. To improve sensitivity while preserving dimensionality, the overall expression was raised to higher powers.

Squaring the expression provided the most accurate correspondence with visually perceived increases in mitochondrial complexity, without altering the underlying information content. The final MCI formulation used for analysis was therefore:

Expanded Dynamic Range Formulation

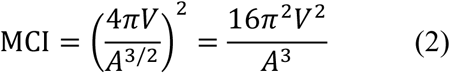

Both equations encode identical structural information. Equation (2) simply expands dynamic range.

### Interpretation of MCI values

- Low MCI values correspond to compact, near-spherical mitochondria with minimal branching.
- High MCI values indicate elongated, branched, or highly complex mitochondrial structures, including networks with multiple junctions or nanotunnels.
- MCI is unbounded, allowing complexity to scale without an upper limit as mitochondrial morphology becomes increasingly elaborate.

### Comparison with sphericity

MCI was evaluated alongside sphericity, a three-dimensional shape descriptor defined as:

Comparison with Sphericity (restated cleanly)

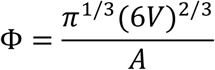

Sphericity is bounded: 0 < Φ ≤ 1

MCI is unbounded.

Sphericity asymptotically approaches 1 for a perfect sphere and decreases with increasing elongation or irregularity. However, because sphericity is bounded between 0 and 1, it saturates at high levels of morphological complexity and becomes less sensitive to differences among highly branched mitochondria. MCI was therefore selected as the primary metric of mitochondrial branching and complexity, as it captures theoretically unlimited increases in structural complexity arising from fusion, network expansion, and nanotunnel formation.

### Statistical handling

MCI values were calculated for each reconstructed mitochondrion, averaged across biological replicates in downstream statistical analyses. All surface-area and volume inputs were derived from the same 3D triangulated surface mesh to ensure internal consistency of the metrics.

### Cell Culture

Human and mouse primary myotubes were isolated and cultured in DMEM/F-12 (Gibco, Waltham, MA, USA) with 20% FBS, 10 ng/mL basic fibroblast growth factor, 1% penicillin/streptomycin, 300 μL/100 mL Fungizone, 1% non-essential amino acids, and 1 mM β-mercaptoethanol. C2C12 myoblasts were maintained in Dulbecco’s modified Eagle’s medium (DMEM) containing 10% fetal bovine serum, 4.5 g/L glucose, and 1% Penicillin-Streptomycin. The media was changed every other day after washing with phosphate-buffered saline (PBS) to remove residual media components.

### Primary Myotube Culture and ATF4 Expression Studies

As previously mentioned, satellite cell isolation was carried out (Pereira, Tadinada et al. 2017, Stephens, Mungai et al. 2023, Vue, Garza-Lopez et al. 2023, Scudese, Marshall et al. 2025). Culture dishes coated with BD Matrigel were used to plate primary satellite cells extracted from Atf4fl/fl mice (n = 8 per experimental condition). The cells were grown in DMEM-F12 supplemented with 20% FBS, 40 ng/mL bFGF, 1× nonessential amino acids, 0.14 mM β-mercaptoethanol, 1× penicillin/streptomycin, and Fungizone. Myoblasts were maintained in a growth medium supplemented with 10 ng/mL bFGF until they reached around 90% confluency, at which point they were transferred to DMEM-F12 with 2% FBS and 1× insulin-transferrin-selenium to stimulate myotube differentiation. Myotubes were infected with an adenovirus expressing GFP-Cre three days following differentiation to produce ATF4 knockout (ATF4 KO). The ATF4 gain-of-function (ATF4 OE) model was achieved by infecting differentiated myotubes with ATF4-expressing adenovirus (Ad5CMV-ATF4/RSVeGFP).

### Myotube Serial Block-Face Scanning Electron Microscopy and 3D Volumetric Analysis

We used serial block-face scanning electron microscopy (SBF-SEM) to look at differentiated ATF4 KO, ATF4 OE, and WT myotubes. Cells were embedded in resin after being fixed according to standard techniques for electron microscopy. Then, an SBF-SEM platform was used to obtain serial image stacks spanning the entire cell volume. These datasets enabled quantitative analysis of myotubes using ultrastructural imaging, three-dimensional reconstruction, and volumetric sampling. We used standard methods for SBF-SEM datasets to do the three-dimensional segmentation and rendering. The mitochondrial volume, surface area, sphericity, cristae structure, and organelle interaction parameters were all included in the quantitative studies, along with other relevant measures. Across all biological replicates, all image capture and analyses were performed blinded.

### RNA Extraction and Real-Time qPCR

The RNeasy Kit (Qiagen Inc.) was used to isolate the total RNA, and a NanoDrop 1000 spectrophotometer (NanoDrop Products, Wilmington, DE, USA) was used to measure absorbance at 260 and 280 nm. We used the High-Capacity cDNA Reverse Transcription Kit (Applied Biosystems, Carlsbad, CA, USA) to reverse transcribe 1 μg of RNA. RT-qPCR was carried out utilizing SYBR Green chemistry (Life Technologies, Carlsbad, CA, USA) in accordance with established protocols (Boudina, Sena et al. 2007). Each experimental condition was conducted in triplicate, using about 50 ng of cDNA per reaction, on 384-well plates. The samples were then analyzed using an ABI Prism 7900HT system (Applied Biosystems). The primer sequences can be found in Supporting Information S1: Table 1 (Supplementary File). The manufacturer’s recommended thermal cycling conditions were used.

### Assessment of Mitochondrial H₂O₂ Levels

To determine the amounts of mitochondrial hydrogen peroxide (H₂O₂) production in the ATF4 OE, ATF4 KO, and control groups, cells were seeded in 35 mm dishes at 0.2 million cells per dish or at a comparable density in multiwell plates. Cells were allowed to adhere overnight. To reduce endoplasmic reticulum stress, cells were stimulated for 72 hours with either a vehicle or 4-phenylbutyric acid (4-PBA; 500 µM).

We used the mitochondrial-targeted fluorescent probe MitoPY1 (5 µM; Bio-Techne) to quantify mitochondrial H₂O₂. The cells were then treated with MitoPY1 and incubated in phenol-red-free DMEM for 45 minutes at 37 °C. Prior to fluorescence measurements, the cells were rinsed twice with 1X HBSS and then maintained in fresh HBSS. Fluorescence was measured utilizing a plate reader with MitoPY1-specific excitation/emission parameters (510 nm excitation, 540-560 nm emission). For each condition, the background fluorescence was eliminated using dye-free controls, and the intensity of MitoPY1 fluorescence was standardized to either cell number or total protein content. The results are presented as the relative amounts of mitochondrial H₂O₂ in comparison to the control group of cells. Every experiment was carried out using a minimum of three separate biological replicates and several technical replicates for each condition.

### Seahorse Extracellular Flux Analysis

Cellular respiration was quantified using a Seahorse XF24 extracellular flux (XF) bioanalyzer. The myoblasts from mice were cultured into myotubes by plating them at a density of 2 × 10⁴ cells per well. Following the three-day differentiation period, myotubes were either ATF4 overexpressed or knocked out using the methods previously mentioned. Seahorse assays were conducted three days following genetic modification. Before analysis, cells were cultured in a non-CO₂ incubator for 60 minutes to allow temperature and pH equilibration, and then culture media was exchanged with XF-DMEM (Agilent, 103680). Basal oxygen consumption rate (OCR) was measured first. Then, we sequentially injected oligomycin (1 μg/mL), carbonyl cyanide 4-(trifluoromethoxy)phenylhydrazone (FCCP; 1 μM), rotenone (1 μM), and antimycin A (10 μM) to evaluate ATP-linked respiration, maximal respiration, and non-mitochondrial respiration, respectively (Pereira, Tadinada et al. 2017).

For glycolytic assays, cells were transferred to glucose-free XF-DMEM and kept in a non-CO₂ incubator for an extra half an hour prior to the glycolysis stress test. Triplicate Seahorse plates were used for our studies; the findings presented here are indicative of three separate biological investigations, each of which included four to six technical replicates for each condition.

### ER Stress Inhibition and FGF21 Secretion in ATF4-Overexpressing Myotubes

Differentiated overexpressing ATF4 myotubes were exposed to either a vehicle or 500 μM 4-phenylbutyric acid (PBA) for three days, starting four days after adenoviral infection to prevent stress endoplasmic reticulum (ER) stress. On the third day of treatment, the culture media was changed to differentiation media (phenol-red-free DMEM/F12 with 2% fetal bovine serum), and new PBA was introduced at 8:00 AM. We collected the conditioned media after 8 hours of incubation to analyze for secreted FGF21. Cells were then extracted for protein analysis by Western blotting and RNA isolation for quantitative PCR (qPCR). Following the manufacturer protocol, conditioned media and corresponding cell lysates were analyzed for FGF21 levels using a BioVendor FGF21 ELISA kit (Asheville, NC, USA)(Pereira, Tadinada et al. 2017). Each experiment had four to six biological replicates per condition and was repeated three times.

### In Vitro Exercise Stimulation

In vitro exercise simulation was accomplished by utilizing electrical pulse excitation, as described earlier (Evers-van Gogh, Alex et al. 2015): Differentiated human myotubes, primary mouse myotubes, or C2C12 cells were subject to serum starvation in FBS-free DMEM prior to stimulation. Media was changed right before stimulation. In C-dish culture systems, carbon electrodes were used to administer electrical stimulation utilizing a C-Pace 100 pulse generator (IonOptix, Milton, MA, USA). The stimulation was set at a frequency of 1 Hz, with a pulse duration of 2 ms and an intensity of 11.5 V, for either 4.5 or 24 hours. Conditioned medium from both stimulated and non-stimulated cells were gathered, centrifuged for five minutes at 800 rpm (17 rcf), and then stored at -80°C. Prior to cell collection and lysis for RNA isolation for qPCR or Western blot studies, the target cell lines were treated with a 1:1 mixture of growth medium containing 10% FBS and conditioned medium that did not contain FBS.

### Pharmacological Induction of Cellular Stress

For chemical induction of cellular stress, C2C12 myoblasts were treated with vehicle control (DMSO), 1 μg/mL tunicamycin or 2 μg/mL thapsigargin for 8 hrs prior to being washed and incubated with staining solution for live-cell imaging.

### Compound 4f and WRR139 Chemical Inhibition Studies

C2C12 myoblasts were treated with control (DMSO), Compound 4f (10 μM for 48 hrs; Nrf2-in-1; MedChemExrpress) for Nrf2 inhibition, WRR139 (10 μM for 1 hour; MedChemExpress) for Nrf1 inhibition, or both Compound 4f (10 μM for 47 hrs prior to WRR139 addition) and WRR139 (10 μM for 1 hr prior to being washed and incubated in staining solution) and incubated following the manufacturer’s guideline prior to cellular stress induction. All inhibitors were maintained in medium throughout cellular stress and imaging studies.

### Live-cell Imaging and Analysis

C2C12 myoblasts were seeding into 35 mm glass-bottom dishes for live-cell imaging studies. A staining solution consisting of phenol and serum free medium, 100 nM MitoTracker Green for mitochondrial morphology and subcellular distribution analysis, and 50 nM tetramethylrhodamine, methyl ester (TMRM) to measure membrane potential as an assessment of mitochondrial function. Cells were washed with sterile PBS and incubated in the staining solution for 30 minutes at 37⁰C. Following incubation, cells were washed twice with sterile PBS prior to the medium being replaced with phenol-free DMEM. Z-stacks of live cells were obtained using a Nikon Ti2-E inverted fluorescence microscope equipped with a Yokogawa CSU-W1 with a spinning disc confocal system, 100X Plan Apo NA 1.45 objective, Hamamatsu Fusion BT camera, piezo stage controller, an environmental chamber, solid-state laser diodes (405, 488, 561, and 640 nm), and SoRa super-resolution module. All components were controlled using NIS-Elements software (version 5.42; Nikon, Melville, NY). Emission filters for the d488 nm channel were 525/36 nm and 605/52 nm for the 561 nm channel. W1 confocal mode was used to acquire serial optical sections with a z-step size of 300 nm.

### Myotube Validation

Myoblasts were seeded and incubated in differentiation medium for 3-7 days. 3-days post-differentiation, myotubes were infected with adenovirus expressing GFP. 48 hrs. post infection, myotubes were fixed in 4% paraformaldehyde (PFA) for 5 minutes prior to being washed 3X with PBS. Cells were then incubated for 10 minutes in permeabilization buffer and then for 1 hour in blocking solution at 22⁰C (Stephens, Mungai et al. 2023). Blocking solution was then aspirated and cells were incubated in a solution of blocking buffer and SC-71-s primary antibody for myosin heavy chain Type IIA overnight at 4⁰C. The following day the primary antibody solution was aspirated, and cells were washed PBS 3X for 5 minutes each, then incubated in a solution of secondary antibody and blocking solution for 1 hour at 22⁰C in in a dark room. Cells were then incubated with DAPI (1 μg/mL) in PBS for 5 minutes. Cells were then washed and mounting media applied. Coverslips were then placed on the slides and given adequate time to dry prior to being sealed and imaged using light microscopy.

### Nucleoid Staining and Imaging

Myoblasts were seeded on 35 mm glass-bottom dishes and either differentiated into myotubes or fixed and images when confluence was reached. Media was aspirated from myoblasts and myotubes and replaced with a working solution of warm media, SYBR Gold nucleic acid gel stain, MitoTracker Orange CMTMRos and incubated for 35 minutes at 37⁰C. The working solution was then aspirated, and cells were washed with warm media. Fixation was performed with 4% PFA and 0.5% glutaraldehyde for 10 minutes at 37⁰C, rinsed 3X with sterile PBS and mounted with Vectasheild and allowed to sufficient curing time before being subject to confocal microscopy.

### Three-Dimensional Reconstruction and Mitochondrial Morphology Analysis

The electron microscopy images were subsequently imported into ImageJ (NIH) for initial processing and alignment. Using specialized segmentation software, three-dimensional reconstruction of mitochondrial networks was conducted to identify mitochondria from serial-section image stacks. Each individual mitochondria were manually traced across serial sections. Mitochondrial parameters were quantified, which include volume (μm³), surface area (μm²), perimeter (μm), sphericity, and mitochondrial branching index (MBI). Circularity was calculated as 4π × (area/perimeter²), where a value of 1.0 represents a perfect circle. Increased circularity reflects a shift toward rounded mitochondria and loss of elongated mitochondrial networks. Statistical analyses and graphical representations were generated using GraphPad Prism (version 9.0) (Garza-Lopez, Vue et al. 2021, Neikirk, Vue et al. 2023, Crabtree, Neikirk et al. 2024, Crabtree, Neikirk et al. 2024, Hinton, Katti et al. 2024, Shao, Killion et al. 2024, Scudese, Marshall et al. 2025).

### Fly Strains and Genetics

Flies were cultured on standard yeast cornmeal agar medium in vials or bottles at 25 °C under a 12-h light/dark cycle. The Mef2-Gal4 driver line (also known as P{GAL4-Mef2.R}3) was used to direct expression of upstream activating sequence (UAS) transgenes specifically in muscles. UAS-mito-GFP (II) (BDSC#8442) was used to visualise mitochondria. For manipulation of ATF4 signalling, RNAi knockdown and overexpression lines targeting ATF4/cryptocephal (crc) were crossed to Mef2-Gal4. The RNAi line was obtained from the Vienna Drosophila RNAi Centre (VDRC) (line #2935; FBti0084038). ATF4/crc overexpression (OE) lines (FlyORF: F004853) were used where indicated. Chromosome designations and additional strain details are available through FlyBase (http://flybase.org). Male and female flies were analysed together, as no sex-dependent differences in mitochondrial morphology were observed in wild-type muscle. The Mef2-Gal4 strain served as the genetic background control for all experiments.

### Fly RNAseq

All fly RNA-sequencing procedures followed those described in earlier publications (Hall et al., 2017; Ponce et al., 2020). Genomic experts from the University of Iowa’s Institute of Human Genetics were able to extract whole RNA from fly populations. RNA was extracted from five biological replicates. These included control flies, flies with ATF4 (cryptocephal; crc) knockdown (ATF4 KD) and flies with ATF4 overexpression (ATF4 OE). The integrity and purity of the RNA were evaluated before the library was prepared. Kallisto (v0.46.2) was used to pseudo-align and quantify sequencing reads. An index was constructed from a FASTA file of annotated transcripts that corresponded to the Drosophila melanogaster genome assembly BDGP6.28. The DESeq2 (v1.30.0) program was used to perform the differential gene expression analysis. Gene lists were created using absolute fold changes greater than 2.0 and expression levels of more than 5 FPKM in control or ATF4-manipulated samples. Bioinformatics tools such as PANTHER (http://pantherdb.org) for Gene Ontology (GO) classification, Ingenuity Pathway Analysis (IPA; QIAGEN) for upstream regulator and network connectivity analyses, and WebGestalt (http://www.webgestalt.org) for enriched KEGG pathways and transcription factor binding site analysis were used to further examine these gene sets for biological function enrichment.

### RNA sample preparation in flies for qRT-PCR

RNA was extracted from indirect flight muscles (IFMs) that were 1 to 2 days old. TRIzol (Sigma-Aldrich) was used to extract IFMs from the bisected thoraces after they were heated to 4°C. Utilizing a tissue grinder, total RNA was extracted using TRIzol reagent (Invitrogen). To produce cDNA, we used SuperScript III (Invitrogen) and reverse-transcribed the PCR reaction. Using the QuantiStudio 6 Flex equipment (Applied Biosystems) and iTAQ Universal Sybergreen (Biorad), 50 ng of cDNA were utilized for each reaction in quantitative RT-PCR. We used the ΔΔCT method to evaluate gene expression, and we normalized relative expression to GAPDH.

### Mitochondrial Staining and Quantification in FLY

As described previously by Katti et al., (Scudese, Marshall et al. 2025), the IFMs were identified by dissecting the thoraces of adult Drosophila that were 2-3 days old in 4% paraformaldehyde (PF). A 1.5-hour agitation-aided fixation in 4% PF was followed by three 15-minute washing with PBSTx (phosphate-buffered saline + 0.3% Triton X-100) on isolated muscles. The mitochondrial-targeted green fluorescent protein (GFP) generated from UAS-mito-GFP under the direction of Mef2-Gal4, mitochondria were detected. The muscles were incubated in 2.5 μg/mL phalloidin-TRITC (Sigma) in PBS for 40 minutes at 25°C to stain the F-actin protein. A Zeiss LSM 780 confocal microscope was used to capture images of stained muscles that had been mounted in Prolong Glass Antifade Mountant with NucBlue (ThermoFisher).

Fluorescence microscopy was used to image muscle fibers in order to quantify mitochondria. As indicated earlier, mitochondria were labeled using either mito-GFPThe images were taken with a magnification of 60× and processed with the help of ImageJ tool. After the images were segmented, mitochondria that extended over three sarcomeres (from the first sarcomere’s Z-disc to the fourth sarcomere’s Z-disc) were chosen for further examination. Using ImageJ, the number of mitochondria present in three sarcomeres was quantified.

### Statistical Analysis

Results are expressed as mean values ± standard error of mean (SEM). If two groups were compared, unpaired Student’s *t-*tests were employed. For more than two groups, one-way or two-way analysis of variance tests, as appropriate, were applied, followed by posttests employing Tukey’s multiple comparisons or Fisher’s protected least significant difference tests. Statistical analyses were performed using the software programs GraphPad Prism and StatPlus software from SAS Institute. The level of significance in all analyses was fixed at p < 0.05, as marked: *p < 0.05, **p < 0.01, ***p < 0.001, ****p < 0.0001.

## Results

### Aging is associated with preserved anthropometrics but impaired functional capacity

Participants who participated in the investigation were not advised to alter their diet or lifestyle practices during the experimental period. Rather, they were instructed to stick to their regular eating habits and daily schedules. This strategy was used to make sure that participants reflected their typical physiological and environmental circumstances, including their regular training or exercise routines. Maintaining stable eating and lifestyle patterns throughout human physiology investigations is a popular strategy for reducing variability in metabolic and physiological measurements caused by abrupt lifestyle changes. Stable dietary habits are essential for reducing the confounding effects associated with metabolic variability, as skeletal muscle metabolism, mitochondrial activity, and exercise adaptations are all influenced by nutritional status and habitual diet (Lexell 1995, Phillips and Winett 2010, Larsson, Degens et al. 2019, Morton, Murphy et al. 2019).

To assess how aging affects physical attributes and functional capacity in our human cohort, participants were categorized by age and sex, then evaluated for anthropometric, functional, and expression parameters (Figure 1). The young and old groups displayed significant differences in chronological age (average age for young women (19 years) and men (19 years) vs. older women (73 years) and men (74 years),respectively, p<0.0001) whereas height remained stable across all groups (1.6m and 1.8m for young/old women and men, respectively) demonstrating that stature is maintained regardless of age and sex (Figure 1A-B). Body mass index (BMI) [median (IQR) for young women was 22.0 kg/m^2^ (21.2-22.9 kg/m^2^), young men 24.6 kg/m^2^ (21.5-26.4 kg/m^2^), old women 24.0 kg/m^2^ (19.6-26.0 kg/m^2^) and old men 25.7 kg/m^2^ (24.0-28.78 kg/m^2^)] and body weight (young women 58 kg (56-59 kg), young men 75 kg (69-82kg), old women 58kg (49-66 kg) and old men 72kg (69-93kg) exhibited minor fluctuations between groups but lacked consistent age-related patterns, indicating that basic anthropometric measurements alone fail to adequately reflect functional aging processes (Figure 1C-D).

**Figure 1.**
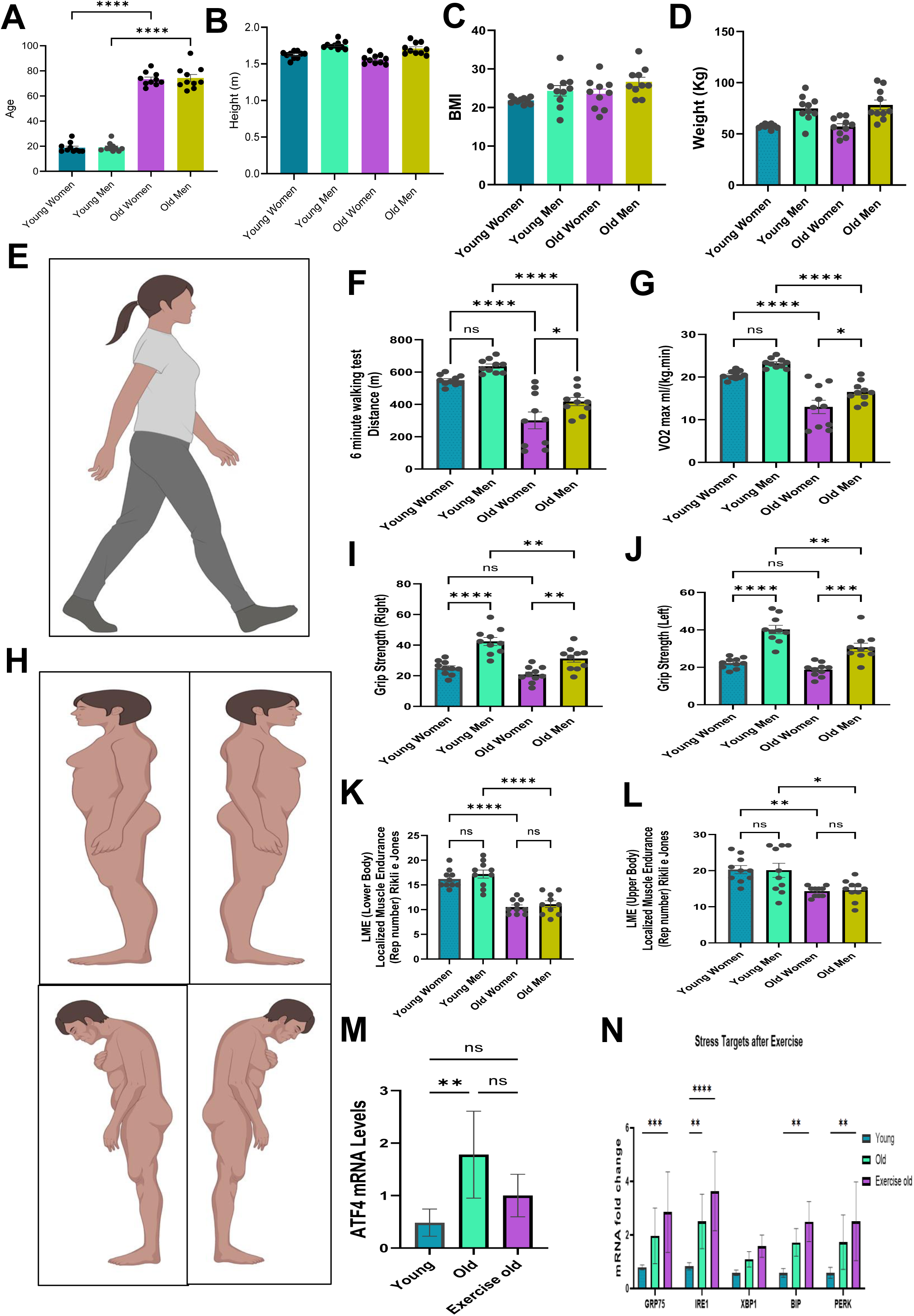
**Sex- and age-dependent differences in physical performance, body composition, and muscle function.** (A–D) Distribution of age (A), height (B), body mass index (BMI; C), and body weight (D) across young and older male and female participant groups. (E) Representative image of the standardized 6-minute walking course used to assess functional aerobic capacity (Created with BioRender.com). (F–G) Quantification of total mean distance walked (F; younger women =549m, younger men =636m, old women =301, old men = 418m) and maximal mean oxygen consumption (VO₂max; G: younger women =20.46 ml/(kg.min), younger men =23.22 ml/(kg.min), old women =13.00 ml/(kg.min), old men = 16.51 ml/(kg.min)) measured at the completion of the 6-minute walking test, stratified by age and sex. (H) Representative images illustrating the assessment of right and left hand grip strength in young and older participants (Created with BioRender.com). (I–J) Comparative quantification of right (I: younger women =25.06, younger men =42.42, old women =20.83, old men = 31.24) and left (J: younger women =22.25, younger men =40.19, old women =18.66, old men = 30.67) hand grip strength across age and sex groups. (K–L) Measures of lower body (K: younger women =16.20, younger men =17.20, old women =10.50, old men = 11.10) and upper body (L: younger women =20.30, younger men =20.10, old women =14.30, old men = 14.60) localized muscle endurance, demonstrating age- and sex-specific declines in neuromuscular performance. (M) ATF4 mRNA expression levels across young (0.484) and old (old =1.78 and old who exercise= 1.00)) and (N) mRNA fold changes compared for young, old and exercising old for mitochondria proteins GRP75 (0.79, 1.96, and 2.85, respectively), IREI (0.82, 2.50, and 3.633, respectively), XBP1 (0.58, 1.09, 1.58, respectively), BIP (0.58, 1.72, 2.49, respectively), CHOP (0.58, 1.78, 2.58, respectively), PERK (0.58, 1.73, 2.51, respectively). All data are presented as mean ± SEM with individual data points overlaid. Young and older cohorts consisted of 10 males and 10 females per age group (total *n* = 40). Statistical comparisons were performed using age × sex two-way analyses, with significance defined as *p* < 0.05.

***Aging is associated with impaired aerobic capacity and mobility performance*** Despite the relative stability in anthropometric values, the older group shows some decline in functional ability. Distance achieved in the six-minute walk test is significantly lower in the older female and older male groups in comparison with the younger groups (Figure 1E-F). Maximal Oxygen Uptake (VO2 max) is significantly lower in the older group than in the younger group (Figure 1G). There is a reduction in VO2 max values irrespective of gender. Interestingly, there was no significant difference in VO2 max between younger males vs. younger females. However, in the older cohorts where older males record higher VO2 max in comparison to older females. Taken together, these results suggest the onset of age-related decline in functional ability despite the lack of significant change to BMI and body weight, which strongly highlights the vital roles of functional variables to the biology of aging.

### Muscle strength declines with age in a sex-dependent manner

To better assess neuromuscular function, grip strength in both hands was also quantified (Figure 1H). Compared to the young, both right and left grip strength in older individuals was lower (Figure 1I-J). While young men possessed superior overall grip strength, aging caused a drastic reduction in both men and women, with older females having the lowest. Muscular endurance in both the lower and upper body, measured using resistance-based functional tests, showed similar impairments with aging. Older subjects performed fewer repetitions in lower body endurance tests and upper body endurance tests than younger subjects (Figure 1K-L). These results suggest that aging is associated with lower muscular strength and endurance capacity in a widespread manner but not specific functions.

### ATF4 expression is elevated with aging and modulated by exercise

Given the age-related decline in physical function, we decided to investigate the relationship between aging, as well as exercise, and ATF4 gene expression in human skeletal muscle. ATF4 gene expression levels were found to be higher in older subjects compared to the young subjects (Figure 1M). Interestingly, the level of ATF4 gene expression in older subjects who exercise was not significantly different from that of the young subjects. These findings suggest that ATF4 is dynamically regulated in the human muscle in response to both aging and exercise.

### Exercise activates ATF4-linked stress response pathways

To establish whether there was corresponding expression of other stress-related genes, we quantified the expression levels of classical ATF4-related genes and integrated stress-response target genes. Exercise was associated with stress-responsive transcripts, such as GRP75, IRE1, XBP1, BiP, and PERK, that were higher in older individuals when compared with the younger control subjects (Figure 1N).

Collectively, these data suggest that older age in humans is associated with marked deficits in aerobic capacity, mobility, strength, and endurance that occur despite relative stability in the underlying anthropometric variables.

### Aging is associated with mitochondrial enlargement and enhanced network fusion in murine quadriceps muscle

To assess the effect of progressive aging on mitochondrial network organization in skeletal muscle, we analyzed the quadriceps of young (3 months old) and aged (2-year-old) mice using three-dimensional quantitative analysis.

Three-dimensional reconstructions of mitochondria with the orthoslices removed, highlight age-related changes in the mitochondrial network (Figure 2 A-B’). Looking at transverse and longitudinal views of the reconstructions, aging was characterized by a prominent tendency toward more elongated and highly interconnected mitochondria with fused networks extending greater distances along the myofibrillar direction (Figure 2 A–A’, B–B’). Taken together, these data suggest that aging tends to induce an increase in size and fusion of the mitochondrial network in mouse quadriceps muscle.

**Figure 2.**
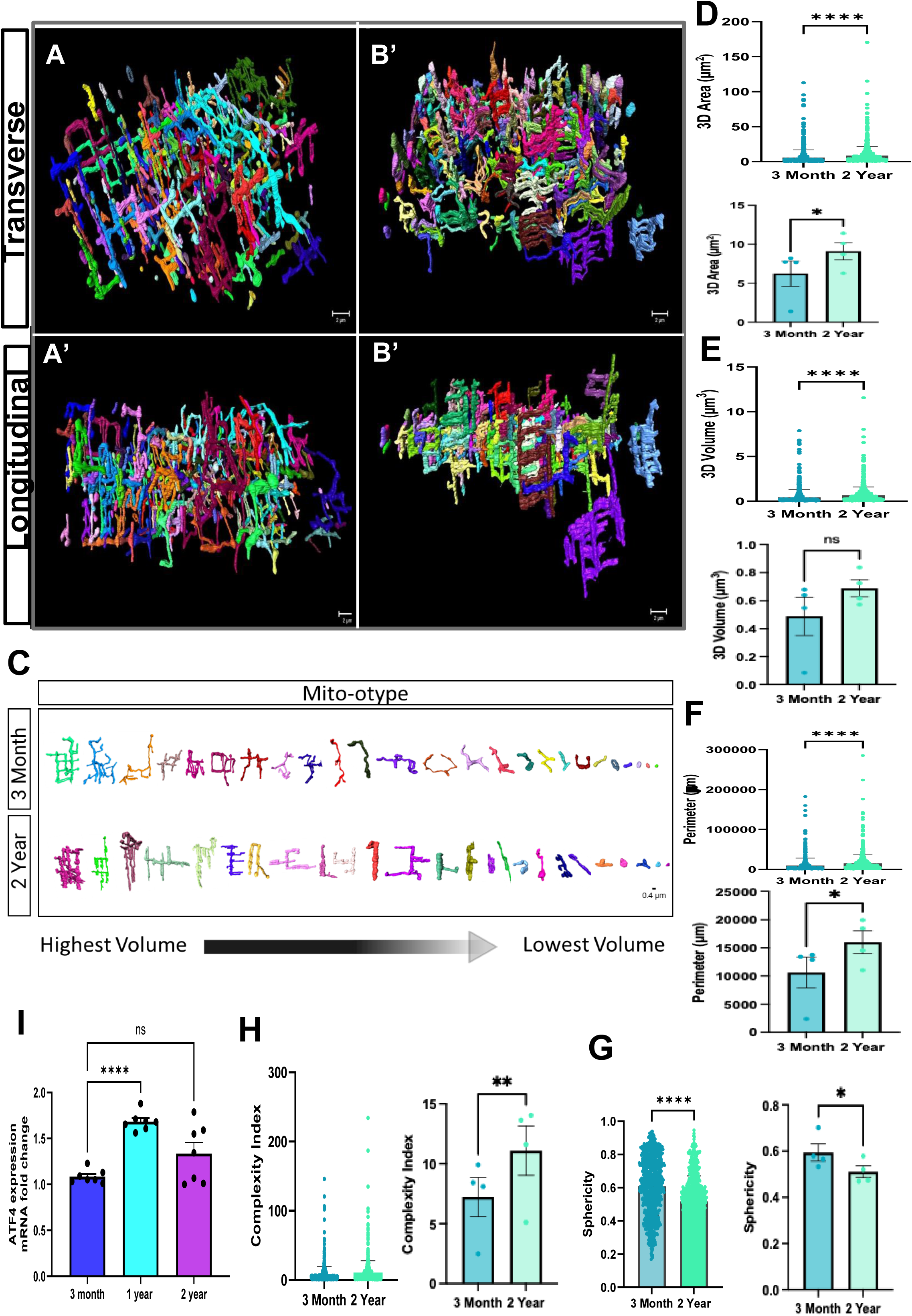
**Age-associated mitochondrial enlargement, fusion, and ATF4 upregulation in murine quadriceps muscle.** (A–B) Representative three-dimensional reconstructions of mitochondria with orthoslices removed, revealing age-associated changes in network organization of quadriceps muscle from 3-month-old (A) and 2-year-old (B) mice in (A–B) Transverse views and (A’–B’) longitudinal views of reconstructed mitochondria, demonstrating a pronounced shift toward larger, more elongated, and fused mitochondrial structures in aged quadriceps muscle. (C) Mitochondrial “mito-typing” based on volume classes, illustrating a redistribution toward higher-volume mitochondrial populations with aging. (D–H) Quantitative 3D morphometric analyses comparing young and aged quadriceps muscle, including mitochondrial area (D), volume (E), perimeter (F), sphericity (G), and mitochondrial complexity index (H). These analyses reveal significant age-dependent increases in mitochondrial size and network complexity consistent with enhanced fusion and remodeling. (I) qPCR analysis of quadriceps muscle demonstrates a significant age-dependent increase in ATF4 expression, in contrast to the gastrocnemius muscle, which does not exhibit ATF4 upregulation with aging. This muscle-specific transcriptional response suggests differential activation of stress-adaptive signaling pathways in weight-bearing versus locomotor muscles. Data are presented as mean ± SEM with individual data points shown. Statistical comparisons were performed using unpaired two-tailed *t* tests. Significance thresholds are indicated as *p* < 0.05, **p** < 0.01, ***p*** < 0.001, **p** < 0.0001. Scale bars, 1 μm.

### Quantitative 3D reconstruction and analysis confirm age-dependent increases in mitochondrial size and complexity

To determine the extent of the changes in the structure of mitochondria with age, thorough three-dimensional morphometry was applied to reconstructed volumes of mitochondria. Mitotyping, as based on size, demonstrated that there was an alteration towards the dominance of higher-volume mitochondria in the ageing quadriceps muscle, when compared with younger tissue (Figure 2 C). Further, aged quadriceps muscle compared to aged muscle had significantly higher three-dimensional area (5.69 vs. 8.78 µm^2^, p<0.0001; Figure 3 D), volume (0.67 vs. 0.43 µm^3^, p<0.0001; Figure 3 E), as well as the perimeter (15476 vs. 96.71 µm, p<0.0001; Figure 3 F) of mitochondria (Figure 2 D-F).

**Figure 3.**
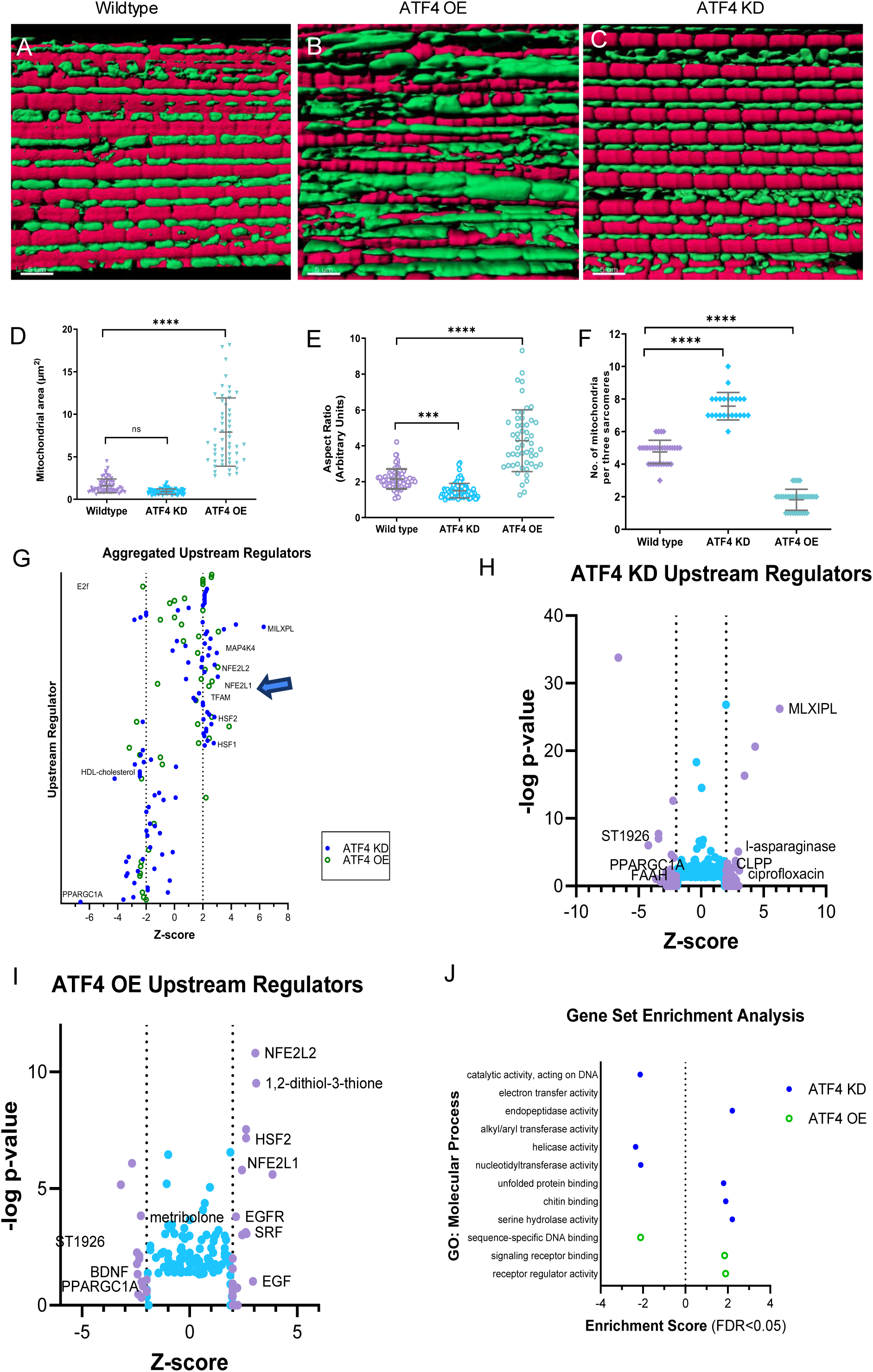
**ATF4 bidirectionally regulates mitochondrial morphology and transcriptional programs controlling mitochondrial organization and biogenesis in muscle.** (A–C) Representative confocal volumetric reconstructions of muscle fibers from wildtype (A), ATF4 overexpression (ATF4 OE; B), and ATF4 knockdown (ATF4 KD; C) models. Actin filaments are rendered in magenta and mitochondria in green. Confocal imaging reveals regularly aligned mitochondrial arrays in wildtype muscle. ATF4 OE induces pronounced mitochondrial enlargement, elongation, and clustering between actin filaments, whereas ATF4 KD results in smaller, fragmented mitochondria with increased organelle density and disrupted alignment. (D–F) Quantitative analysis of confocal-derived mitochondrial morphology, including mitochondrial area (D), aspect ratio (E), and number of mitochondria per sarcomere (F). ATF4 OE significantly increases mitochondrial size and elongation while reducing mitochondrial number per sarcomere, consistent with enhanced mitochondrial fusion. In contrast, ATF4 KD reduces mitochondrial size and elongation and increases mitochondrial number, consistent with mitochondrial fragmentation. (G) Aggregated upstream regulator analysis from RNA-sequencing comparing ATF4 KD and ATF4 OE muscle, plotted by activation Z-score. TFAM is identified as a prominent ATF4-responsive regulator. (H–I) Predicted upstream regulators in ATF4 KD (H) and ATF4 OE (I) muscle. ATF4 KD suppresses mitochondrial and metabolic transcriptional networks, whereas ATF4 OE activates stress-adaptive and mitochondrial regulatory pathways converging on TFAM, NRF2 (NFE2L2), and growth-associated signaling. (J) Gene set enrichment analysis (GSEA) highlighting molecular function categories significantly altered by ATF4 manipulation. ATF4 KD preferentially downregulates transcriptional and mitochondrial maintenance pathways, whereas ATF4 OE enriches signaling and regulatory programs associated with mitochondrial remodeling. Data are shown with individual measurements overlaid. Statistical significance is denoted as ns, *p* < 0.001, p < 0.0001. Scale bars, 5 μm.

In keeping with these size changes, the sphericity index of mitochondria was lower in 2-year mice compared to 3-month mice (0.52 vs. 0.61, p<0.0001), implying an alteration from spherical mitochondria towards more elongated, merged types (Figure 2 G). In addition, the complexity index, indicative of connectivity, was higher in older muscle mitochondria compared to the young (10.51 vs. 6.89, p<0.0001), implying that branching patterns were enhanced (Figure 2 H). These three quantitative observations combined suggest that mitochondrial enlargement, surface area, complexity, and branching ability all increase during aging in the mouse quadriceps muscle.

### Mitochondrial morphology changes are accompanied by muscle-specific upregulation of ATF4

Given the importance of ATF4 in mediating cellular stress responses and directing adaptive change in mitochondria, we next investigated how ATF4 expression might change with age in skeletal muscle. Using qPCR, we found that ATF4 mRNA expression was significantly higher in the 1-year-old group than in the young 3-month group (Figure 2I). However, this increase was not observed in the 2-year age group. Interestingly, these findings indicate that although there were reductions in ATF4 mRNA expression with aging in mouse gastrocnemius samples, there were no changes in ATF4 mRNA expression observed in the 2-year age group in the quadriceps samples, despite an increase in the 1-year age group.

### ATF4 Remodeling Is Associated with Changes in Mitochondrial Architecture in *Drosophila* Indirect flight muscle as revealed by confocal volumetric imaging

We analyzed mitochondrial morphology in cells with altered ATF4 expression to better understand ATF4’s role in regulating mitochondrial structure and cellular stress responses. The wild-type, ATF4 overexpression (ATF4 OE), and ATF4 knockdown (ATF4 KD) conditions were clearly distinguished by high-resolution imaging. Wild-type cells have an ordered mitochondrial network with elongated structures throughout the cytoplasm. By contrast, when ATF4 was overexpressed, mitochondrial architecture was significantly remodeled, with increased mitochondrial number, altered aspect ratio, and increased mitochondrial area. The quantitative study of mitochondrial morphology revealed that ATF4 KD cells differ significantly from ATF4 OE cells, indicating that ATF4 plays a role in controlling mitochondrial dynamics and organelle organization. According to these results, structural alterations in the mitochondrial network are linked to ATF4 activity, indicating that transcriptional stress responses impact mitochondrial architecture.

To address the question of whether ATF4 directly affects mitochondrial morphology in vivo, we analyzed the organization of the mitochondria within the Indirect flight muscle of ATF4 KD or ATF4 OE *Drosophila* relative to wild-type controls using confocal volumetric microscopic imaging (Figure 3 A-C). Analysis of the confocal reconstructions revealed significant genotype-dependent differences in mitochondrial morphology. Within the wild-type muscle tissue, the mitochondria were arrayed in a regular fashion between the myofibrils that were elongated (Figure 3 A). Overexpression of ATF4 caused significant mitochondrial elongation with the formation of clusters of these structures with loss of normal sarcomeric organization (Figure 3 B). Smaller punctate mitochondria were observed upon ATF4 KD with increased density but reduced elongation, suggestive of a fragmented mitochondrial network (Figure 3 C).

Mitochondria are elongated and form a clustered network and fragmented mitochondria. Quantitative analysis of the mitochondrial morphology from the confocal images confirmed these results. Over-expression of ATF4 strongly led to an increase in the mitochondrial area (wildtype vs ATF4 OE, 1.58 vs. 7.91, p<0.0001; wildtype vs ATF4 KD, 1.58 vs. 0.91, p=0.087; Figure 3 D) and the mitochondrial aspect ratio (wildtype vs ATF4 OE, 2.15 vs. 4.28, p<0.0001; wildtype vs ATF4 KD, 2.15 vs. 1.50 Figure 3 E) but to a reduction in the mitochondrial number per sarcomere (wildtype vs ATF4 OE, 4.75 vs. 1.81, p<0.0001; wildtype vs ATF4 KD, 4.75 vs. 7.57 Figure 3 F), which could indicate mitochondrial fusion events. On the other hand, mitochondrial aspect ratio diminished, while the number of mitochondria per sarcomere increased with ATF4 KD, which could indicate mitochondrial fission events (Figure 3 D-F). These results indicate the sufficiency and necessity of ATF4 for regulating mitochondrial morphology.

### ATF4-dependent mitochondrial remodeling is linked to transcriptional control of TFAM-centered networks in Drosophila flight muscle

To clarify the transcriptional mechanisms underlying ATF4-induced mitochondrial restructuring, we performed RNA sequencing and further analyzed upstream regulators and gene sets. The aggregated upstream regulator result showed clear differences between ATF4 KD and ATF4 OE, with opposite patterns of activation in the regulation of mitochondria and metabolism (Figure 3G). Notably, a key ATF4-targeting upstream regulator was TFAM, which corresponds to ATF4’s role in integrating the restructuring of mitochondria with the transcriptional regulation of gene expression in mitochondria.

ATF4 KD muscle also presented the upstream regulators showing the repression of TFAM-related pathways and the downregulation of transcriptional regulation of mitochondrial function (Figure 3 H). ATF4 OE triggered activation of pathways related to TFAM and stress-response transcription factors like Nrf2, as well as cell-growth pathways (Figure 3I). Gene set enrichment further validated these observations, showing that ATF4 KD mainly suppressed the regulation of DNA binding and transcription regulation, whereas ATF4 OE mainly enriched the pathways associated with mitochondrial remodeling (Figure 3 J).

Genomic sequencing was carried out to explore deeper into transcriptional pathways linked to ATF4 disruption. Regulatory pathways involved in proteostasis, mitochondrial function, and cellular stress responses were more abundant among the differentially expressed genes. Upstream regulator analysis identified several transcription factors thought to be responsible for the observed transcriptional alterations. Among these, Nuclear Factor Erythroid 2 Like 1 (NFE2L1), also known as Nrf1, appeared as a prominent predicted upstream regulator. Nrf1 belongs to the transcription factor family known as the cap’n’collar basic leucine zipper (CNC-bZIP). This family is defined by a basic leucine zipper motif that is essential for dimerization and DNA binding, as well as a conserved CNC domain. Multiple physiological stressors, such as oxidative stress, metabolic stress, and proteostasis disruptions, activate Nrf1, which is expressed ubiquitously throughout tissues. Numerous physiological functions are controlled by this transcription factor, including the defense against oxidative stress, inflammatory signaling, metabolism, cell differentiation, and protein homeostasis. Therefore, the RNA-seq data imply that regulatory mechanisms in stress adaption and proteostasis interact with ATF4-dependent transcriptional processes.

Taken together, these data establish that ATF4 affects mitochondrial morphology as a function of transcriptional programs that ultimately target TFAM, providing insight into the transcription-mediated adaptation of the mitochondrial network structure. Confocal volumetric imaging indicates that differential changes in the expression levels of various genes influenced by ATF4 affect mitochondrial size, elongation, and localization, suggesting that ATF4 is an essential downstream regulatory element controlling mitochondrial structure in skeletal muscle.

### Mitochondrial nucleoid organization, mitochondrial structure, and ATF4–TFAM activation in primary mouse myotubes and myoblasts

To characterize mitochondrial organization and the ATF4–TFAM axis in muscle cells, we first visualized the architecture of differentiated, multinucleated myotubes, which exhibited nuclear alignment along the longitudinal axis (Figure 4 A), and confirmed their identity using Myosin Heavy Chain Type-IIa immunofluorescence (Figure 4B).

**Figure 4.**
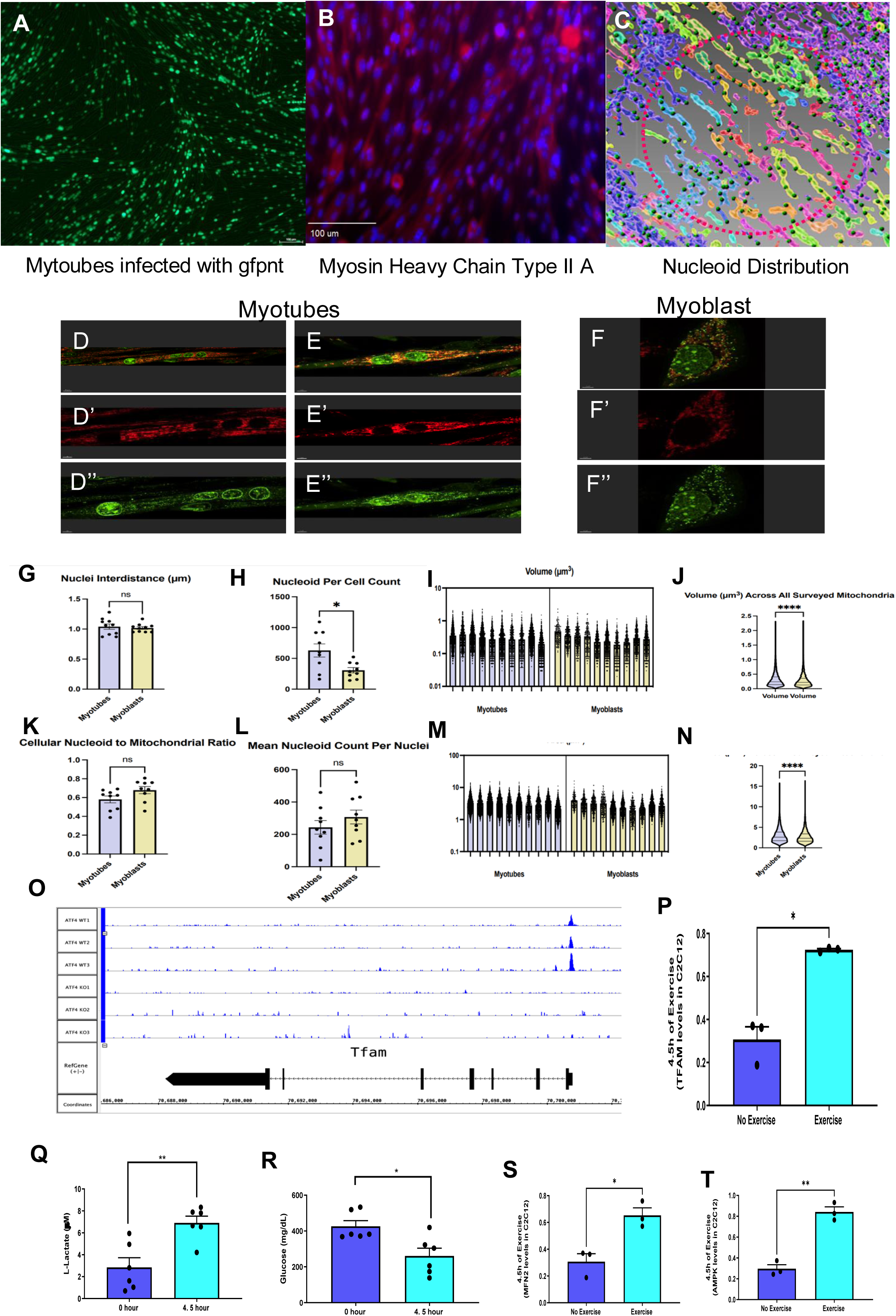
**Mitochondrial nucleoid organization, mitochondrial structure, and ATF4–TFAM activation in myotubes and myoblasts** (A) Representative fluorescence image of differentiated myotubes infected with nuclear-targeted GFP (GFP-NT), highlighting multinucleated myotube architecture and nuclear alignment along the longitudinal axis. (B) Immunofluorescence staining for Myosin Heavy Chain Type IIa, confirming myotube differentiation and fiber-type identity. (C) Computational segmentation and spatial mapping of mitochondrial nucleoid distribution within myotubes. Dashed outlines indicate individual myotubes, revealing that nucleoid distribution is not always uniform, with regions of clustering and spacing heterogeneity along the myotube length. (D–D″) Representative confocal images of myotubes. (D) Merged image showing mitochondria and nucleoids. (D′) MitoTracker staining showing mitochondrial networks alone. (D″) SYBR Gold staining highlights mitochondrial nucleoids. (E–E″) Independent myotube examples shown in the same format as (D–D″), demonstrating reproducibility of mitochondrial and nucleoid organization patterns. (E′) MitoTracker staining. (E″) SYBR Gold nucleoid staining. (F–F″) Representative confocal images of myoblasts shown in the same channel separation as myotubes. (F) Merged image. (F′) MitoTracker staining of mitochondria. (F″) SYBR Gold staining of mitochondrial nucleoids. (G) Quantification of inter-nucleoid distance (µm) in myotubes versus myoblasts. (H) Nucleoid number per cell, revealing differences in nucleoid abundance between differentiated and undifferentiated muscle cells. (I) Distribution of mitochondrial volume (µm³) across individual mitochondria in myotubes and myoblasts. (J) Violin plot summarizing mitochondrial volume (µm³) across all surveyed mitochondria. (K) Cellular nucleoid-to-mitochondrial ratio comparing myotubes and myoblasts. (L) Mean nucleoid count per nucleus, indicating scaling of mitochondrial genetic content with nuclear number. (M) Distribution of mitochondrial area (µm²) in myotubes and myoblasts. (N) Violin plot summarizing mitochondrial area (µm²) across all analyzed mitochondria. (O) Integrated visualization of ATF4 chromatin occupancy across the *Tfam* genomic locus, supporting direct transcriptional regulation of *Tfam* by ATF4. (P) Acute in vitro exercise-like stimulation of myotubes (e.g., electrical pulse stimulation or metabolic contraction paradigm) significantly increases *Tfam* expression relative to unstimulated controls, demonstrating activation of the ATF4–TFAM axis under contraction-like conditions. (Q–R) Metabolic responses to exercise-like stimulation in myotubes, including (Q) Increased extracellular lactate accumulation, and (R) Reduced glucose levels in the culture medium, consistent with enhanced metabolic demand. (S–T) Activation of mitochondrial remodeling and energy-sensing pathways in stimulated myotubes, including (S) Induction of Mitofusin 2 (MFN2), consistent with enhanced mitochondrial fusion and organelle connectivity, and (T) Activation of AMPK, indicating engagement of energy-sensing and metabolic stress signaling pathways. Data are presented as mean ± SEM with individual data points shown. Statistical significance is denoted as ns (not significant), *p* < 0.05, p < 0.01, *p* < 0.001, p < 0.0001. Statistical analyses were performed using unpaired two-tailed *t* tests or one-way or two-way ANOVA with appropriate post hoc testing, as indicated.

Computational segmentation of mitochondrial nucleoids within myotubes revealed a non-uniform spatial distribution, with heterogeneous regions of clustering and spacing along the myotube length (Figure 4 C). Representative confocal imaging of myotubes showed organized mitochondrial networks and nucleoids (Figure 4 D–D″), a pattern reproducible in independent examples (Figure 4 E–E″). In contrast, myoblasts displayed distinct mitochondrial and nucleoid organization (Figure 4 F–F″). Quantification revealed that inter-nucleoid distance was not significantly different in myotubes versus myoblasts (1.04 vs. 1.02, p=0.650; Figure 4 G), but nucleoid number per cell differed and was significantly higher in myotubes vs. myoblasts (600 vs 289, p=0.0244; Figure 4 H). Analysis of individual mitochondrial morphology showed distributions of mitochondrial volume (Figure 4 I and Figure 4 J) and area (Figure 4 M and Figure 4 N) in both cell types. The cellular nucleoid-to-mitochondrial ratio (Figure 4 K) and the mean nucleoid count per nucleus (Figure 4 L) were also assessed, indicating scaling of mitochondrial genetic content.

Molecular analysis demonstrated direct transcriptional regulation, as integrated visualization showed ATF4 chromatin occupancy across the *Tfam* genomic locus (Figure 4 O). Furthermore, compared to no exercise, acute exercise-like stimulation of myotubes significantly increased *Tfam* expression (0.30 no exercise vs. 0.72 acute exercise, p=0.0192; Figure 4 P). This stimulation also triggered metabolic responses, including increased extracellular lactate accumulation (2.85 vs 6.91 µM, p=0.0045; Figure 4 Q) and reduced culture medium glucose levels (426 vs 261mg/dl, p=0.0130; Figure 4 R). Finally, stimulated myotubes exhibited activation of mitochondrial remodeling and energy-sensing pathways, with induction of Mitofusin 2 (MFN2) (0.31 vs. 0.65, p=0.0135; Figure 4 S) and activation of AMPK (0.29 vs. 0.84, p=0.0012; Figure 4 T).

Studies have demonstrated that asymmetric cell division can occur, making TFAM distribution during cell division of interest. We hypothesize that enhanced TFAM expression after ATF4 overexpression may cause mitochondrial nucleoid organization alterations. Mitochondrial nucleoids seem to be distributed quite uniformly in myoblasts, although they become more dispersed in myotubes. ATF4 overexpression may further induce unequal nucleoid clustering or structure. Further evidence that mitochondrial nucleoid structure may vary dynamically in reaction to physiological stress is the observation that TFAM levels seem to rise after exercise. Aging may exacerbate these effects, especially in myotubes with diminished differentiation capacity.

### TFAM and mtDNA Maintenance Link Stress Signaling to Mitochondrial Regulation

The mitochondrial remodeling regulation. Transcription factor A (TFAM) is essential for the transcription, replication, and organization of mitochondrial DNA, and it is also an essential component of mitochondrial DNA maintenance. The number of mitochondrial DNA copies and its structural stability are both regulated by TFAM. Mutations or loss of TFAM function impair oxidative phosphorylation and interfere with mitochondrial gene expression. Nuclear Respiratory Factor 1 (NRF1) is a transcription factor that regulates the expression of TFAM in numerous cellular systems. These systems are involved in respiratory chain assembly and mitochondrial biogenesis. Research has shown that ATF4 signaling modulates NRF1-TFAM signaling, which in turn affects mitochondrial regulatory pathways. Research has demonstrated that during stress, ATF4 activation inhibits NRF1 expression, which in turn lowers TFAM levels and makes mitochondrial DNA maintenance less efficient. These findings lay the groundwork for a potential mechanism by which stress-induced ATF4-mediated reactions can alter mitochondria in our experimental setup.

### ATF4 is sufficient and necessary to regulate mitochondrial size and network architecture in primary cells and skeletal muscles

To directly examine whether ATF4 regulates mitochondrial morphology and positioning, we analyzed the ultrastructure of mitochondria in primary cells derived from three different groups of mice: controls, ATF4 KO, and ATF4 OE. We analyzed SBF-SEM micrographs of these cells using three-dimensional reconstructions. Orthoslices obtained from these images revealed clear differences among the genotypes in mitochondrial positioning (Figure 5A-C). While those of controls were of intermediate sizes with moderate elongation and a fair degree of connectivity between adjacent ones, those of ATF4 KO cells tended to be of a smaller size with less elongation and a lesser degree of connectivity. Indeed, the contrary was true of ATF4 OE cells.

**Figure 5.**
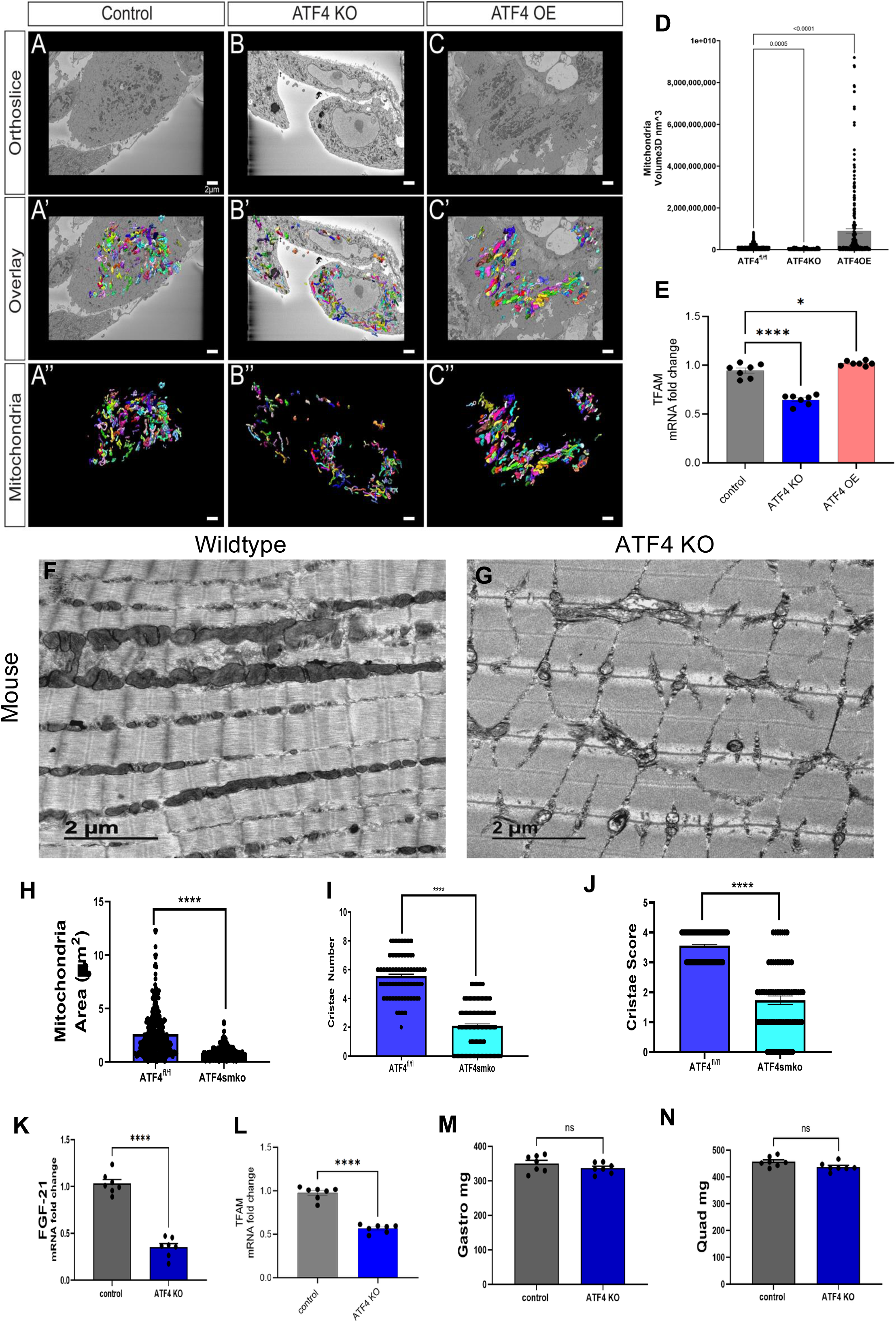
**ATF4 bidirectionally regulates mitochondrial morphology, network organization, and cristae architecture in primary cells and skeletal muscle.** (A–C) Representative transmission electron microscopy (TEM) orthoslices from primary cells isolated from control (A), ATF4 knockout (ATF4 KO; B), and ATF4 overexpression (ATF4 OE; C) mouse models. (A′–C′) Corresponding orthoslices with three-dimensional (3D) reconstructed mitochondria overlaid, illustrating genotype-dependent differences in mitochondrial distribution, size, and spatial organization within the cellular volume. (A″–C″) Isolated 3D mitochondrial reconstructions with orthoslices computationally removed, revealing simplified and fragmented mitochondrial networks following ATF4 deletion and enlarged, elongated, and more interconnected mitochondrial structures following ATF4 overexpression. (D) Quantitative 3D morphometric analyses of mitochondria volume (D). ATF4 deletion results in significant reductions in mitochondrial volume and (E) TFAM expression levels, whereas ATF4 overexpression significantly increases mitochondrial volume, consistent with enhanced mitochondrial elongation and fusion. (F–G) Representative TEM images from skeletal muscle of wild-type (F) and skeletal muscle–specific ATF4 knockout (ATF4 smKO; G) mice. Wild-type muscle exhibits elongated, well-aligned mitochondria with preserved internal structure, whereas ATF4-deficient muscle displays fragmented, disorganized mitochondria with altered spatial arrangement relative to myofibrils. (H–J) Quantitative analyses of mitochondrial ultrastructure in skeletal muscle, including mitochondrial area (G), cristae number (I), and cristae score (J). ATF4 deficiency results in significant reductions in mitochondrial size and marked impairment of cristae organization, indicating compromised inner membrane architecture. (K) FGF-21 and (L) TFAM mRNA fold change was significantly reduced in knockout (KO) compared to control models while (M) Gastrocnemius (Gastro) and (N) Quadriceps (Quad) muscle mass (mg) between control and knockout (KO) models were similar. Data are presented as violin plots or bar graphs with individual data points shown, and median and interquartile ranges or mean ± SEM indicated as appropriate. Statistical comparisons were performed using one-way ANOVA with post hoc testing or unpaired two-tailed t tests, as indicated. Significance thresholds are denoted as ns (not significant), *p* < 0.05, **p** < 0.01, ***p*** < 0.001, and **p** < 0.0001. Scale bars, 2 μm (A–C) and 2 μm (G–H).

Overlaying reconstructed mitochondria on orthoslices helped to further define the discrepancies in the distribution and patterns of mitochondrial organization within the cell volume (Figure 5 A′–C′). Digital processing of three-dimensional reconstructed mitochondria showed that ATF4 deficiency results in less complex networks with reduced elongation and branchingmitochondrial structures, whereas ATF4 overexpression results in larger, more interconnected complexes (Figure 5 A″–C″).

### Quantitative 3D morphometric analyses reveal bidirectional ATF4 control of mitochondrial size and complexity

To establish the impact of ATF4 modulation on mitochondrial morphology, we analyzed cells with ATF4 KO and OE using quantitative three-dimensional analysis to assess the robustness of our findings. Cells with ATF4 knockout showed substantial lower mitochondrial volume, while cells with ATF4 overexpression showed substantially higher mitochondrial volume (Figure 5 D). Moreover, TFAM mRNA expression is increased in the ATF4 OE group and decreased in the ATF4 KO group, indicating an association with ATF4 (Figure 5E) (Casas-Martinez, Xia et al. 2025).

### ATF4 is required for maintenance of mitochondrial size and cristae organization in skeletal muscle in vivo

To determine whether the ATF4-mediated mitochondrial remodeling we detected in primary cells is maintained in living organisms, we analyzed the ultrastructure of mitochondria in the skeletal muscles of wild-type mice and those in which ATF4 is deleted.

While skeletal muscles in wild-type mice demonstrated normally stretched. Well-oriented mitochondria along the axes of the myofibers, as depicted in Figure 5 F, those in ATF4-deleted mice were highly disoriented and appeared to be smaller in size, as depicted in Figure 5 G. This was shown by quantitative measurements to be true, as there was a significant decrease in the mitochondrial area within the muscle tissue of ATF4 knockout mice compared to wild-type mice (Figure 5 H). Moreover, the number and score of cristae were significantly reduced in muscle tissue from ATF4 knockout mice (Figures 5 I-J). These results make it clear that ATF4 regulating mitochondrial volume and global architecture but also in regulating cristae architecture within skeletal muscle. The mRNA quantitative measurements showed that FGF21 and TFAM expression decreased in the ATF4 KO compared to the control group (Figure 5 K-L). The quantitative analysis for gastro and quadriceps muscle showed decreased weight in the ATF4 KO group (Figure 5 M-N).

Taken together, these results show that ATF4 is a bidirectional mediator of mitochondrial shape and ultrastructure. The strong correlation between primary cell-based systems and in vivo skeletal muscle suggests that ATF4 is a regulator of mitochondrial shape in both stress conditions and physiological states.

### ATF4 regulates mitochondrial respiratory capacity and bioenergetic flexibility in myotubes ATF4 directly engages the Tfam locus

To determine whether ATF4 directly regulates the transcription of Tfam, we analyzed the ATF4 chromatin occupancy across the Tfam locus. Analysis of the genome browser showed in the Tfam regulatory region, which was reduced in the absence of ATF4 (Figure 6 A). This data further confirms direct transcriptional linkages between ATF4 and Tfam, thereby placing TFAM downstream of ATF4 signaling.

**Figure 6.**
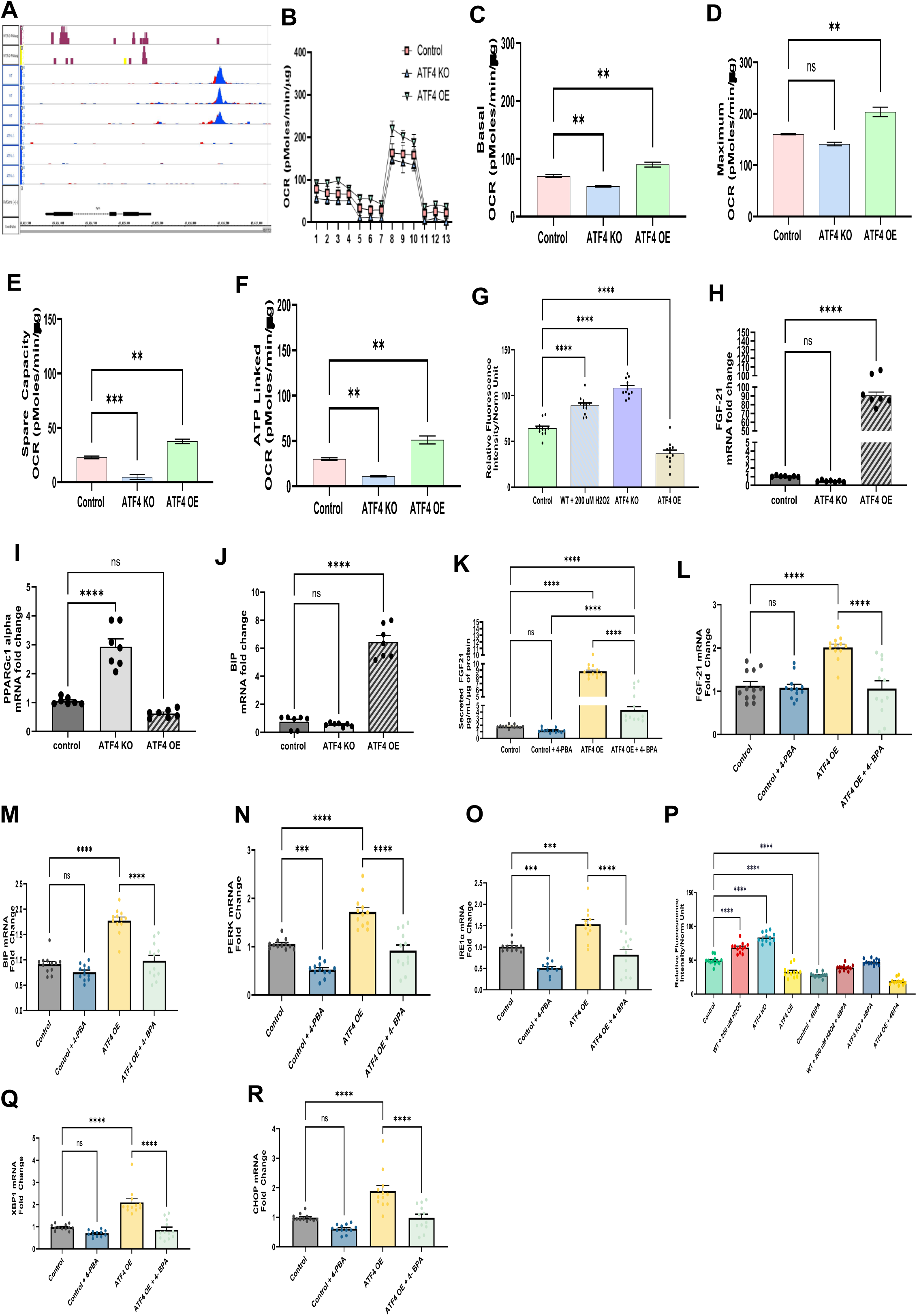
**ATF4 integrates mitochondrial bioenergetics, redox homeostasis, and TFAM-dependent transcriptional adaptation during in vitro exercise–like stimulation in myotubes.** (A) Genome browser tracks showing ATF4 chromatin occupancy across regulatory regions of the *Tfam* locus under control, ATF4 knockout (ATF4 KO), and ATF4 overexpression (ATF4 OE) conditions, indicating ATF4-dependent transcriptional regulation of *Tfam*. (B) Seahorse extracellular flux OCR time course following sequential mitochondrial stressors in differentiated myotubes under control, ATF4 KO, and ATF4 OE conditions. (C–F) Quantification of mitochondrial respiration parameters in myotubes, including basal OCR (C), maximal respiration (D), spare respiratory capacity (E), and ATP-linked OCR (F). ATF4 KO significantly impairs mitochondrial oxidative capacity, whereas ATF4 OE enhances respiratory performance and bioenergetic flexibility. (G) Relative fluorescence–based measurement of reactive oxygen species (ROS) in myotubes under basal conditions and following oxidative challenge. ATF4 KO myotubes exhibit elevated ROS accumulation, while ATF4 OE myotubes demonstrate improved redox control. (H–K) Quantitative PCR (qPCR) analysis of stress-response and mitochondrial regulatory genes in myotubes. ATF4 OE induces transcriptional programs associated with mitochondrial adaptation and proteostasis, whereas ATF4 KO suppresses these responses, including reduced *FGF-21* expression (H), *Tfam* expression (I), increased *PPARGc1-alpha* (J) and reduced *BIP expression*. (L–P) Pharmacologic modulation of metabolic and stress signaling pathways in myotubes reveals ATF4-dependent regulation of mitochondrial and stress-response genes, including reduced (L) secretion of FGF21, significant changes in mRNA expression fold change in (M) *FGF-21, (N) Bip*, (O) *PERK,* (P) IRE1-alpha, (Q) XBP1, (R) CHOP, and (T) *Tfam*. These changes are seen with relative fluorescence intensity across different groups. Data are presented as mean ± SEM with individual data points shown. Statistical significance is denoted as ns (not significant), *p* < 0.05, p < 0.01, *p* < 0.001, p < 0.0001. Statistical comparisons were performed using one-way or two-way ANOVA with appropriate post hoc testing or unpaired two-tailed *t* tests, as indicated.

To understand the function of ATF4 in mitochondrial bioenergetics in a setting of heightened metabolic demand, we evaluated mitochondrial respiration in differentiated myotubes using Seahorse extracellular flux analysis. Wild-type, ATF4 KO, and ATF4 OE myotubes differed dramatically in OCR traces in response to a series of mitochondrial stressors (Figure 6 B).

Quantitative analysis showed that basal OCR values were lower in ATF4 KO myotubes but higher in ATF4 OE myotubes than in controls (Figure 6 C). Maximal respiratory capacity was equally attenuated in ATF4 KO myotubes but augmented in ATF4 OE myotubes (Figure 6 D). In line with these findings, spare respiratory capacity, an indicator of mitochondrial ability to accommodate an elevated energy requirement, was attenuated in ATF4 KO but augmented in ATF4 OE myotubes (Figure 6 E). These data indicate that ATP-linked respiratory capacity, a measure of mitochondrial ability to support ATP production during a stress condition, was increased with forced ATF4 expression (Figure 6 F). Taken together, these findings demonstrate ATF4 to be an important regulator of the oxidative capacity of mitochondria in myotubes.

### ATF4 limits mitochondrial ROS accumulation and promotes redox resilience during metabolic stress

As alterations in mitochondrial respiration are intimately linked to reactive oxygen species (ROS), we proceeded to assess redox status in cells within myotubes with altered ATF4 levels. In contrast, those ROS levels than those in normal cells (Figure 6). Using fluorescence-based ROS measurements, we confirmed that basal ROS levels were elevated in ATF4 KO myotubes, whereas those in ATF4 OE were lower than in normal cells (Figure 6G). Following oxidative stress, ATF4 KO cells compared with control cells; on the other hand, ATF4 OE cells exhibited superior redox capacity. Collectively, these data indicate that ATF4 has a role in potentiating both mitochondrial respiration and redox capacity to counter oxidative stress when cells are exposed to heightened metabolism.

#### ATF4 controls transcriptional stress responses and mitochondrial regulatory programs, including TFAM

To transcript levels of stress-response and mitochondrial regulatory genes in myotubes. Quantitative PCR analysis indicated a significant induction of stress-response transcripts in ATF4 OE myotubes, while there was significant suppression in the ATF4 KO myotubes (Figure 6 H-J).

Specifically, a critical regulator of mitochondrial transcription, Tfam, was decreased in ATF4 KO myotubes but increased in ATF4 OE myotubes compared to respective controls (Figure 6 I). This transcriptional response was not limited to mitochondrial transcriptional regulators but also included other stress-responsive genes, such as Bip and Chop, among others known to be targets of ATF4.

Pharmacological modulation of metabolic and stress pathways further showed that ATF4-dependent induction of Tfam and related genes is indeed driven by cellular stress signals, imparting even greater importance to ATF4’s function as an integrator of metabolism and mitochondrial transcriptional responses (Figure 6 K-O). The profiling of ATF4-dependent stress and metabolic gene XBP1, CHOP1, and IRE1-alpha decreases in the 4-PBA group and increases ATF4 OE myotubes, demonstrating coordinated regulation of mitochondrial, redox, and unfolded protein response pathways (Figure 6 P-R).

### Pharmacologic treatment of ATF4 signaling alters mitochondrial spatial organization under basal and ER stress conditions in C2C12 myoblasts

We investigated the function of the NGLY1-Nrf1 pathway to to stress-adaptive transcriptional responses. WRR139 is a small chemical that pharmacologically inhibits cytosolic peptide:N-glycanase (NGLY1), an enzyme necessary for Nrf1 to be properly activated during proteasome stress. The endoplasmic reticulum is responsible for N-glycosylation during normal cellular synthesis of Nrf1. Following retrotranslocation to the cytosol, the protein is degraded via the ER-associated degradation pathway. Due to NGLY1-dependent deglycosylation and proteolytic processing, Nrf1 can evade total degradation in the presence of reduced proteasome activity. This procedure produces an active transcription factor that can enter the nucleus and activate transcription of proteasome subunit genes. The proteasome “bounce-back” reaction, which repairs proteasome capacity after inhibition, is comprised of this process. This pathway is disrupted when NGLY1 is inhibited with WRR139. Blocking NGLY1 activity prevents Nrf1 from processing correctly and from activating proteasome gene transcription. Consequently, proteotoxic stress accumulates within the cell, and adaptive recovery of proteasome function is inhibited. Our RNA-seq results show transcriptional alterations, but these findings provide a bigger picture of disrupted proteostasis signaling.

To investigate the impact of ATF4 signaling on mitochondrial spatial arrangement, mitochondrial fluorescence intensity was quantified across defined concentric cellular zones (nuclear, perinuclear, central, radial, and distal) under basal conditions and following pharmacological induction of endoplasmic reticulum (ER) stress. Experimental groups included cells treated with vehicle (DMSO), the ATF4 pathway modulator WRR139, the ATF4-inhibiting compound 4f, or their combination, both in the absence and presence of thapsigargin or tunicamycin (Figure 7 A–R).

**Figure 7.**
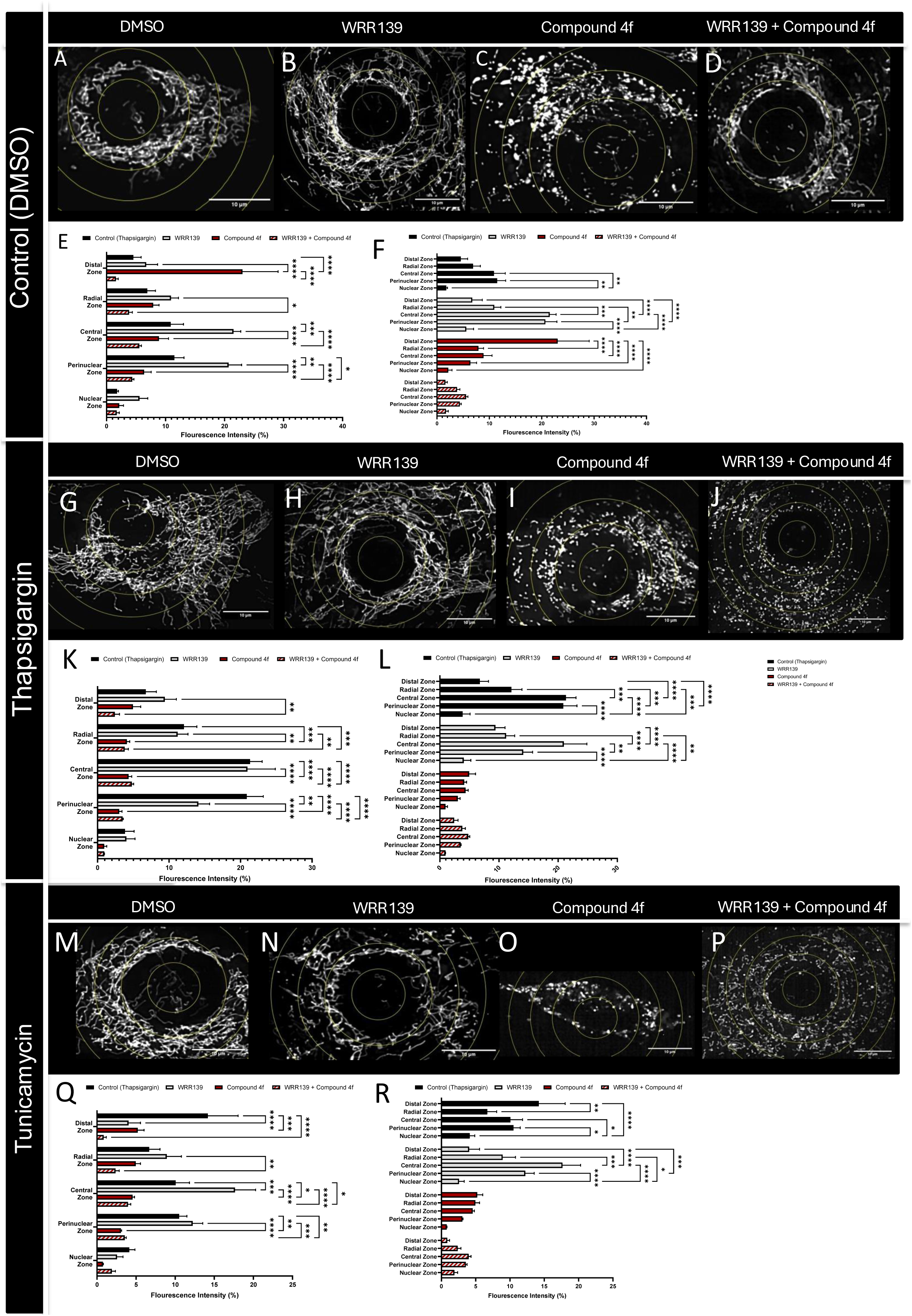
**Spatial redistribution of mitochondrial networks under ER stress and pharmacologic modulation** (A–D) Representative high-resolution images of mitochondrial networks in cells treated with DMSO (control), WRR139, Compound 4f, or WRR139 + Compound 4f under basal conditions. Concentric radial masks (yellow) define nuclear, perinuclear, central, radial, and distal zones used for spatial quantification of mitochondrial signal intensity. (E–F) Quantification of mitochondrial fluorescence intensity (%) across defined radial zones under control (DMSO) conditions. Bar plots summarize redistribution of mitochondrial signal following treatment with WRR139, Compound 4f, or their combination, revealing treatment-dependent alterations in mitochondrial positioning relative to the nucleus. (G–J) Representative mitochondrial network images following thapsigargin-induced ER stress, shown for DMSO, WRR139, Compound 4f, and WRR139 + Compound 4f conditions. ER stress induces pronounced remodeling of mitochondrial organization, including redistribution toward central and perinuclear compartments. (K–L) Quantification of mitochondrial fluorescence intensity (%) across radial zones under thapsigargin treatment, demonstrating that pharmacologic modulation significantly alters stress-induced mitochondrial redistribution patterns. (M–P) Representative mitochondrial network images following tunicamycin-induced ER stress, shown for DMSO, WRR139, Compound 4f, and WRR139 + Compound 4f conditions. Tunicamycin elicits distinct mitochondrial spatial remodeling compared with thapsigargin, indicating stress-specific mitochondrial responses. (Q–R) Quantification of mitochondrial fluorescence intensity (%) across radial zones under tunicamycin treatment, revealing compound-specific effects on mitochondrial positioning and spatial organization during ER stress. Data are presented as mean ± SEM. Statistical significance is denoted as *p* < 0.05, p < 0.01, *p* < 0.001, p < 0.0001. Statistical comparisons were performed using one-way or two-way ANOVA with appropriate post hoc testing, as indicated

In the presence of DMSO (control), mitochondria demonstrated a characteristic concentration within perinuclear and central cellular regions, with signal intensity gradually diminishing toward peripheral areas (Figure 7 A, E–F). Administration of WRR139 alone resulted in a moderate redistribution of mitochondrial signal away from the perinuclear area toward more peripheral regions, indicative of partial initiation of stress-adaptive remodeling pathways. Conversely, treatment with compound 4f significantly reduced overall mitochondrial fluorescence intensity across all regions, with the most substantial decreases occurring in the perinuclear and central zones, suggesting disruption of mitochondrial maintenance or biogenesis. Importantly, combined treatment with WRR139 and compound 4f partially restored mitochondrial signal distribution, yielding a more uniform spatial profile but with drastically blunted fluorescence intensity.

#### ATF4 signaling regulates mitochondrial redistribution during thapsigargin-induced ER stress

Thapsigargin-induced ER stress resulted in pronounced ATF4-dependent mitochondrial remodeling. In cells exposed solely to thapsigargin, there was a significant increase in mitochondrial fluorescence intensity within the perinuclear and central zones, accompanied by a relative reduction in distal regions (Figure 7 G, K–L). This pattern is consistent with stress-induced clustering of mitochondria near the nucleus, characteristic of adaptive integrated stress response (ISR) activation (Kim, Kim et al. 2014).

Co-administration of WRR139 during thapsigargin treatment similarly promoted mitochondrial concentration within the perinuclear and central zones (Figure 7 H, K-L). Conversely, treatment with compound 4f markedly diminished this effect, resulting in decreased perinuclear mitochondrial accumulation and an overall reduction in mitochondrial signal (Figure 7 I, K-L). Quantitative analysis demonstrated significant declines in fluorescence intensity throughout all zones, supporting the notion that Nrf2 may be required for ATF4-mediated maintenance of mitochondrial integrity and spatial organization under acute ER stress. Notably, combined treatment with WRR139 and 4f combined showed similar results to 4f treatment alone with induced cellular stress.

### ATF4 Is Essential for Mitochondrial Maintenance During Tunicamycin-Induced Proteotoxic Stress

To determine whether ATF4-dependent remodeling occurs in other ER stress contexts, mitochondrial organization following tunicamycin treatment was evaluated. Analogous to the SERCA pump inhibitor, thapsigargin, tunicamycin, an inducer of the unfolded protein response, led to a marked redistribution of mitochondria toward perinuclear and central zones, in addition to the distal zone in control cells (Figure 7 M, Q–R), consistent with ISR activation. WRR139 amplified this adaptive mitochondrial clustering (Figure 7 N, Q-R), while compound 4f again elicited substantial reductions in mitochondrial fluorescence intensity (Figure 7 O, Q-R), especially within perinuclear regions. The extent of mitochondrial depletion observed under tunicamycin mirrored that seen with thapsigargin, suggesting that Nrf2 is required for ATF4-mediated mitochondrial spatial stability across different modalities of ER stress. Co-treatment with WRR139 and compound 4f showed effects similar to those observed with 4f treatment alone (Figure 7 P, Q-R).

### Quantitative zonal analysis confirms ATF4-dependent regulation of mitochondrial spatial integrity

Quantitative zonal analysis across all experimental conditions confirmed that ATF4 activity serves as a primary regulator of mitochondrial spatial resilience. Induction of ER stress fostered ATF4-dependent enrichment of mitochondria in the perinuclear and central zones, whereas pharmacological inhibition of Nrf2 disrupted these adaptive distributions, leading to global mitochondrial loss and spatial disarray.

These findings suggest a context in which disruptions in proteostasis signaling interact with mitochondrial regulatory mechanisms. Prolonged proteotoxic stress results from blocking the NGLY1-Nrf1 pathway, processes. The transcriptional pathways linked to the integrated stress response, including ATF4-dependent gene expression, are activated in these stress conditions. In turn, ATF4 signaling affects the NRF1-TFAM pathway, which in turn affects mitochondrial regulatory networks and possibly changes mitochondrial DNA maintenance. This integrated stress-adaptation model may explain the mitochondrial structural changes seen following ATF4 perturbation.

## Discussion

Skeletal muscle aging is characterized by gradual reductions in metabolic flexibility, strength, and endurance that are disproportionate to alterations in gross anthropometric variables or body mass. Current research continues to establish ATF4 as a key regulator of skeletal muscle bioenergetics in response to stress, mitochondrial structural adaptation with aging, and other processes (Cartee, Hepple et al. 2016). Our results, which combine human cohort studies, animal models,3-D mitochondrial reconstructions, ATF4 genetic manipulation, and both exercise and exercise-like metabolic stimulation, demonstrate ATF4 as a context and muscle-type-dependent regulator of aging, mitochondrial architecture, and adaptive stress signaling.

Age-stratified comparisons confirmed that functional impairments in aerobic capacity, mobility, strength, and muscular endurance occur despite stable BMI, highlighting that static anthropometrics poorly reflect functional aging. Notably, ATF4 expression was elevated in aged human muscle but normalized with exercise, and exercise activated a suite of ATF4-linked ER stress and ISR pathway genes (GRP75, IRE1, XBP1, BiP,, PERK), suggesting ATF4 is engaged in compensatory stress-adaptive systems to maintain muscle function against diminishing physiological reserve. These findings are consistent with previous reports that ATF4 activity increases in aging skeletal muscle and is necessary for age-related muscle atrophy (Ebert, Monteys et al. 2010, Ebert, Dyle et al. 2012, Bongers, Fox et al. 2013). These findings provide significant insight into possible reasons behind aging-related functional impairments being frequently masked by changes in body composition or lifestyle habits.

In our human studies, older people showed significant reductions in aerobic ability, mobility, strength, and muscular endurance despite having comparatively stable BMI and body weight. These results support the idea that static anthropometric measures do not adequately represent functional aging. Crucially, ATF4 may be involved in compensatory stress-adaptive systems that try to maintain muscle function in the face of diminishing physiological reserve, as evidenced by the increased expression of ATF4 in aged human muscle and its normalization with exercise (Miller, Marcotte et al. 2023, Von Ruff, Miller et al. 2025). Further evidence that ATF4 plays a role in coordinating adaptive responses to increased energetic demand in aged muscle comes from exercise-induced activation of the ER stress and ATF4-linked integrated stress response (ISR) pathways (Memme, Sanfrancesco et al. 2023). Furthermore, new evidence points to cellular stress signaling pathways as a potential co-factor in skeletal muscle mitochondrial remodeling. An example would be the activation of ATF4, which is involved in integrated stress response signaling, by the loss or deletion of OPA1, a critical protein for mitochondrial inner-membrane fusion. Mitochondrial structural changes can affect organelle communication and stress-response pathways, as ATF4 activation increases mitochondria–ER tethering. Thus, disruptions in the levels of mitochondrial dynamics proteins, including OPA1, MFN2, and MICOS, may change mitochondrial architecture and stress-response processing in skeletal muscle cells (Hinton, Katti et al. 2024). These data indicate that aging-related muscle changes involve fiber type, metabolism, mitochondrial structure, and inter-organelle communication networks.

Critically, our data shows an interesting paradox: ATF4 expression is enhanced in some elderly muscles which is reduced by exercise, while it is decreased in others. We suggest that this reflects context-dependent regulation of ATF4, wherein mechanical load, metabolic demand, or exercise re-engage ATF4-dependent programs whereas basal aging may decrease adaptive stress signals in some muscle. The muscle-specific upregulation of ATF4 in aged quadriceps, coupled with enhanced mitochondrial size in that same tissue, supports the model that ATF4 facilitates compensatory, load-sensitive adaptation.

In our mouse models, aging induced discordant changes, increasing body weight while reducing mass across multiple hindlimb muscles (quadriceps, plantaris, soleus, EDL, gastrocnemius), indicative of sarcopenic remodeling. High-resolution 3D ultrastructural analysis revealed profound but muscle-specific age-related mitochondrial remodeling. Contrary to a universal model of fragmentation, aging quadriceps exhibited mitochondrial enlargement and increased network complexity, suggesting compensatory adaptation in weight-bearing muscles (Araujo, Vargas et al. 2024, Springer-Sapp, Ogbara et al. 2025). Conversely, other muscles showed signs of fragmentation. This muscle-specificity extended to ATF4 expression, which was upregulated in aged quadriceps but not gastrocnemius, indicating differential engagement of stress-adaptive signaling.

Critically, we demonstrate that ATF4 is both necessary and sufficient to bidirectionally regulate mitochondrial morphology (Xu, Liu et al. 2023, Qureshi, Kim et al. 2025). Loss of ATF4 in vivo and in vitro resulted in reduced mitochondrial size, fragmentation, and significant disruption of cristae architecture. Conversely, ATF4 overexpression promoted mitochondrial elongation, enlargement, and enhanced network fusion. The cristae disorganization in ATF4-deficient muscle reveals a novel role for ATF4 in maintaining inner mitochondrial membrane integrity, providing a possible structural link to bioenergetic dysfunction (Streeter, Persaud et al. 2024). These morphological changes were consistent with ATF4’s bidirectional control over mitochondrial bioenergetics, as ATF4 KO impaired while OE enhanced basal respiration, maximal capacity, and spare respiratory capacity in myotubes.

Our findings indicate that TFAM is differentially regulated by ATF4, with both loss and overexpression of ATF4 eliciting distinct effects on TFAM expression. In the knockout model, ATF4 expression is reduced, while in the overexpression model, ATF4 levels are elevated. Research revealed that mitochondrial transcription factor A (TFAM) is a crucial molecular link between stress signaling and mitochondrial genome preservation since it is required for mitochondrial DNA transcription, replication, and nucleoid packaging. Because TFAM expression is directly related to mitochondrial DNA copy number and mitochondrial biogenesis, modulating ATF4-TFAM signaling provides a potential mechanism for proteostasis stress to modify mitochondrial function (Ameri and Harris 2008). Mechanistically, transcriptomic and chromatin occupancy analyses identified TFAM as a key downstream effector. ATF4 binds to regulatory regions of the Tfam locus, and its manipulation bidirectionally regulates TFAM-centered transcriptional networks governing mitochondrial biogenesis and stress adaptation (Hao, Zhong et al. 2021). While ATF4 OE stimulates pathways related to mitochondrial biogenesis, stress adaptation, and redox regulation, ATF4 deficiency decreases TFAM-centered mitochondrial transcriptional networks (Hao, Zhong et al. 2021). These transcriptional changes are reflected functionally: ATF4 promotes both baseline and maximal respiration, spare respiratory capacity, ATP-linked oxygen consumption rate (OCR), and redox stability. Conversely, the absence of ATF4 leads to bioenergetic rigidity and increased reactive oxygen species (ROS) formation. Together, these findings suggest that ATF4 coordinates transcriptional regulation with ultrastructural remodeling to preserve mitochondrial metabolism under stress, placing it at the junction of mitochondrial structure and function.

By maintaining ATF4 activation and changing TFAM-dependent mitochondrial mechanism, such as blockage of the NGLY1–Nrf1 proteostasis pathway may indirectly influence mitochondrial regulatory networks. According to this combined concept, there is a larger adaptive network that includes proteostasis stress, ISR signaling, and regulation of the mitochondrial genome. During cellular stress, eIF2α phosphorylation halts global protein synthesis to reduce the protein-folding burden of the already stressed ER while selectively increasing ATF4 translation. Disruptions in proteostasis are associated with transcriptional regulation through Nuclear Factor Erythroid 2 Like 1 (Nrf1), a key regulator of proteasome-associated gene expression. Activation of this pathway depends on N-glycanase 1 (NGLY1), which facilitates proper activation of Nrf1 and supports proteasomal recovery response. Disruption of this axis impairs protein degradation which sustains proteotoxic stress. These stress pathways are progressively recognized to intersect with the regulation of mitochondria, remarkably through mitochondrial transcription factor A (TFAM), which oversees mitochondrial DNA maintenance and is regulated by Nuclear Respiratory Factor 1 (NRF1), a known master regulator of mitochondrial biogenesis (Rozpedek, Pytel et al. 2016, Tomlin, Gerling-Driessen et al. 2017, Yoshida, Asahina et al. 2021).

Using pharmacologic, morphological, and computational methods, we show that Nrf2, but not Nrf1, is required for ATF4-mediated mitochondrial changes.. In our pharmacological studies, we found that inhibition of Nrf2 with compound 4f in C2C12 myoblasts suppresses . In contrast,downstream effectors of ATF4 signaling, leading to mitochondrial fragmentation, whereas inhibition of Nrf1 with WRR139 maintains network continuity but alters spatial distribution. Combined treatment produced effects similar to those treated with 4f alone, indicating stress adaptation alone cannot preserve mitochondrial subcellular distribution and structure when the downstream effectors of ATF4 pathways are compromised. Aging causes denervation of fast-twitch muscle fibers as some of the motor neurons that supply them deteriorate. The loss and subsequent reinnervation of motor neurons are a key mechanisms underlying changes in fiber composition. Surviving motor neurons, particularly those associated with slow-twitch motor units, can reinnervate these fibers, causing them to acquire slow-twitch fiber properties. The contractile characteristics, metabolic activity, and cellular architecture of skeletal muscle are all altered as a result of this remodeling processes. Thereby, mitochondrial function and integrity within aging muscle fibers become increasingly important as oxidative metabolism becomes more dependent on it during aging (Hepple and Rice 2016). These alterations are strongly related to mitochondrial composition and function. The mitochondria of slow-twitch fibers are more densely packed and produce ATP through oxidative phosphorylation, allowing them to remain active for longer. However, aging is impaired mitochondrial quality control. These alterations can decrease ATP production and lead to muscular weakness and tiredness (Johnson, Robinson et al. 2013, Hood, Memme et al. 2019).

Furthermore, our current study sheds light on the interplay between proteostasis pathways and mitochondrial homeostasis by offering a more comprehensive framework. The stability of proteins is crucial for cell survival, and when this stability is disturbed, transcriptional responses are triggered that restore a balanced ratio of protein synthesis, folding, and degradation. One of the key regulators of this reaction is Nrf1. Unlike Nrf2, Nrf1 is regulated by ER-associated degradation and post-translational modifications. Newly produced Nrf1 enters the ER membrane, becomes N-glycosylated, and is retrotranslocated for proteasomal degradation. However, when proteasome activity is inhibited, Nrf1 manages to evade degradation by undergoing deglycosylation and proteolytic processing before entering the nucleus and activating proteasome gene expression (Kim, Han et al. 2016).

This process is dependent on the cytosolic enzyme N-glycanase 1 (NGLY1), which degrades N-glycans from retrotranslocated Nrf1. By preventing the proteasome recovery program from activation, inhibition of NGLY1, as we accomplished in this study pharacologically with WRR139, limits correct processing of Nrf1. and in turn, increases the cytotoxicity of proteasome inhibition. We found that disrupting the NGLY1–Nrf1 axis may indirectly affect mitochondrial function by maintaining cellular stress signaling and increasing ATF4-dependent transcriptional responses (Tomlin, Gerling-Driessen et al. 2017, Iyer, Mast et al. 2019). Consistent with this, we found mitochondrial structural alterations in our live-cell imaging studies that are compatible with remodeling of mitochondrial networks mediated by ATF4.

While this study provides significant insights into to molecular mechanisms governing ATF4 in aging skeletal muscle, there are a few limitations that should be noted. First, the human muscle cohort is small; therefore, it will be helpful and necessary to confirm these findings with larger and more diverse groups in future investigations. Additionally, the mechanistic experiments were mostly performed in mice and in vitro model systems, which are convenient but may not adequately represent human physiology. Finally, our data suggests that ATF4 regulates mitochondrial transcriptional programs, including TFAM, although more research is needed to understand the downstream regulatory network and ATF4 signaling dynamics.

In summary, in the current study, we show that ATF4 controls mitochondrial morphology and function through TFAM-centered transcriptional pathways. We used comprehensive approaches to show that ATF4 integrates age-related stress signals, exercise stimuli, and metabolic demand, and its role in skeletal muscle aging is dependent on the tissue environment. Our findings show that ATF4 governs cristae architecture, bioenergetic capacity, and spatial resilience position it as a master regulator of mitochondrial health in muscle. These insights highlight the ATF4-TFAM axis as a potential therapeutic target for preserving mitochondrial integrity and muscle function during aging, suggesting that strategies to modulate this pathway could enhance skeletal muscle resilience in the elderly.

## Data availability statement

All data generated or analyzed during this study are included in this published article and its Supplementary information files. Additional data can be requested from the corresponding author.

## Acknowledgments

UNCF/Bristol-Myers Squibb E.E. Just Faculty Fund, Career Award at the Scientific Interface (CASI Award) from the Burroughs Welcome Fund (BWF) ID # 1021868.01, BWF Ad-hoc Award, NIH Small Research Pilot Subaward 5R25HL106365-12 from the National Institutes of Health PRIDE Program, DK020593, Vanderbilt Diabetes and Research Training Center for DRTC Alzheimer’s Disease Pilot & Feasibility Program, CZI Science Diversity Leadership grant number 2022-253529 from the Chan Zuckerberg Initiative DAF, an advised fund of the Silicon Valley Community Foundation to A.H.J. NSF NRT grant 19-22697 (to A.C.), NSF BPE grant 22-17621 (to A.C.), and the Asness Family Fund through the Frist Center for Autism & Innovation (K. Stassun, PI) (to A.C.). We thank Dr. Christopher M. Adams for providing the Ad5CMVATF4/RSVeGFP and ATF4 floxed mice. We would like to also thank George R. Marcotte for his assistance in optimizing the Ad5CMVATF4/RSVeGFP for myotubes.

## Supplementary Figure

**Figure 1. Age-dependent changes in skeletal muscle mass, mitochondrial morphology, and stress-response gene expression in mouse skeletal muscle.**

(A–F) Quantitative comparison of body weight (BW; A) and individual skeletal muscle weights, including quadriceps (B), plantaris (C), soleus (D), extensor digitorum longus (EDL; E), and gastrocnemius (F), between young (3-month-old) and aged (2-year-old) mice.

(G–I) Representative electron microscopy orthoslices from 3-month (G), 1-year (H), and 2-year (I) mouse quadriceps muscle, with corresponding 3D reconstructed mitochondria overlaid (G′, H′, I′). (G″–I″) Full 3D mitochondrial reconstructions with orthoslices removed. (G‴–I‴) Transverse views and (G⁗–I⁗) longitudinal views of reconstructed mitochondria, illustrating progressive age-associated fragmentation and loss of network complexity.

(J) Distribution of mitochondrial shape and volume classes across age groups. (K–O) Quantitative analyses of mitochondrial ultrastructure, including mitochondrial volume (K), surface area (L), perimeter (M), mitochondrial branching index (MBI; N), and sphericity (O).

(P) Quantitative PCR (qPCR) analysis of skeletal muscle revealing age-dependent downregulation of mitochondrial and stress-response genes, including ATF4, in aged muscle relative to young controls, consistent with declining adaptive stress signaling during aging.

All quantitative data are presented as mean ± SEM with individual data points overlaid. Statistical significance was assessed using age-based comparisons, with *p* < 0.05 considered significant. Scale bars, 1 μm.

