## Supplementary material for "ATF4 Coordinates Transcriptomic and Structural Adaptations in Aging Muscle": Supplemetary File

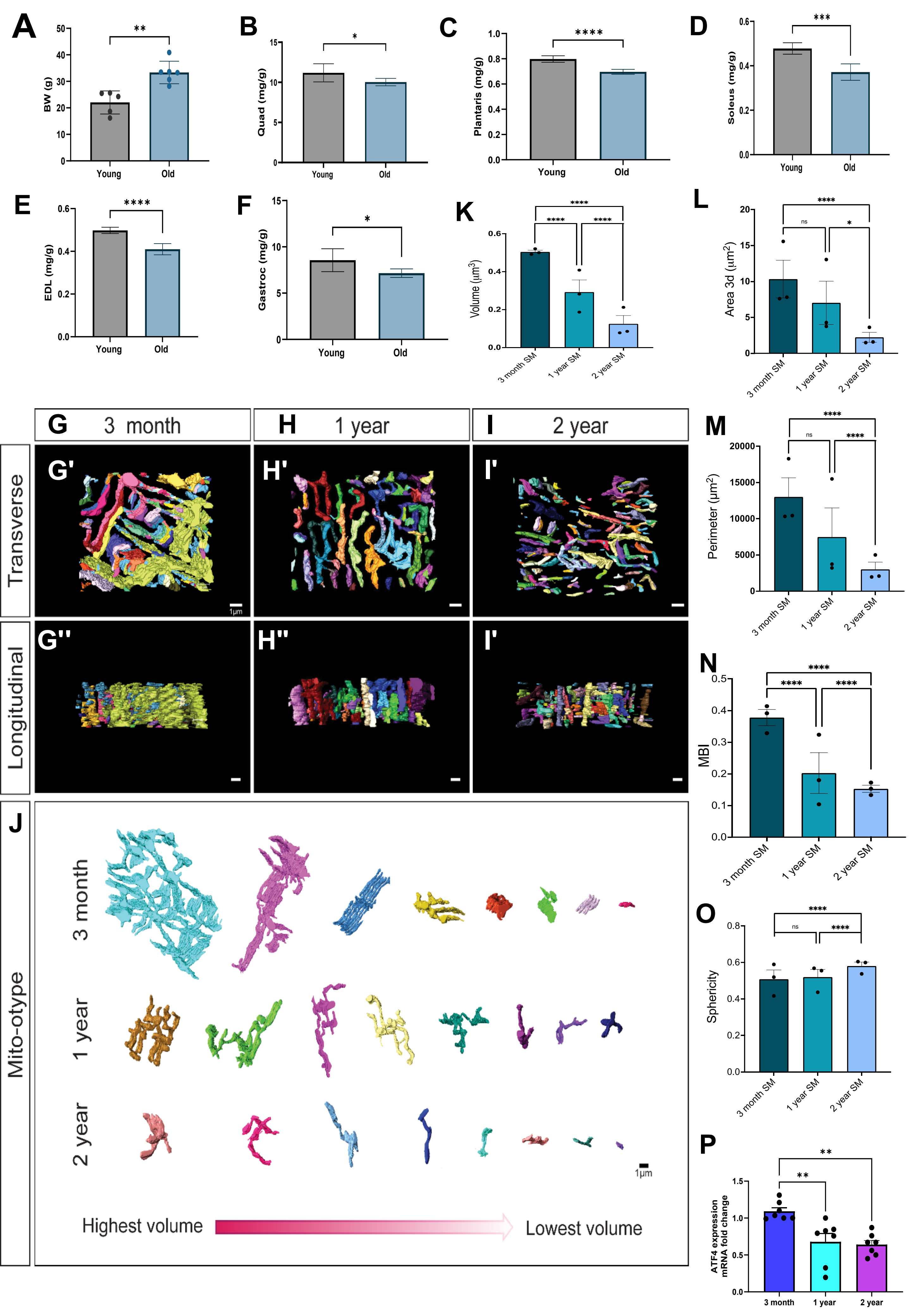

A
BW (g)
Young
Old
**
B
Quad (mg/g)
*
C
Plantaris (mg/g)
****
D
Soleus (mg/g)
***
E
EDL (mg/g)
****
F
Gastroc (mg/g)
*
K
Volume (µm³)
3 month SM
1 year SM
2 year SM
L
Area 3d (µm²)
ns
G
3 month
H
1 year
I
2 year
Transverse
Longitudinal
G'
H'
I'
G''
H''
1µm
M
Perimeter (µm²)
N
MBI
J
Mito-otype
3 month
1 year
2 year
Highest volume
Lowest volume
O
Sphericity
P
ATF4 expression mRNA fold change
3 month
1 year
2 year

### qPCR Primers Used

Human

| Gene | Primers | Sequence |
|---|---|---|
| ***ATF4*** | Forward | 5'- TTCTCCAGCGACAAGGCTAAGG -3' |
| | Reverse | 5'- CTCCAACATCCAATCTGTCCCG-3' |
| ***GPR75*** | Forward | 5'- CCATCAACCTCTCCACTGCCAA-3' |
| | Reverse | 5'- CCATTGCTGGAGAGAACCACCT-3' |
| ***IRE1*** | Forward | 5'- CCGAACGTGATCCGCTACTTCT-3' |
| | Reverse | 5'- CGCAAAGTCCTTCTGCTCCACA-3' |
| ***XBP1*** | Forward | 5'- CTGCCAGAGATCGAAAGAAGGC-3' |
| | Reverse | 5'- CTCCTGGTTCTCAACTACAAGGC-3' |
| ***BIP*** | Forward | 5'-TGTCTTCTCAGCATCAAGCAAGG-3' |
| | Reverse | 5'-CCAACACTTCCTGGACAGGCTT-3' |
| ***PERK*** | Forward | 5'-GTCCCAAGGCTTTGGAATCTGTC-3' |
| | Reverse | 5'-CCTACCAAGACAGGAGTTCTGG-3' |

Mouse

| Gene | Primers | Sequence |
|---|---|---|
| ***FGF21*** | Forward | 5'- ATCAGGGAGGATGGAACAGTGG-3' |
| | Reverse | 5'- AGCTCCATCTGGCTGTTGGCAA-3' |
| ***PPARGc1 alpha*** | Forward | 5'- GAATCAAGCCACTACAGACACCG -3' |
| | Reverse | 5'- CATCCCTCTTGAGCCTTTCGTG -3' |
| ***BIP*** | Forward | 5'- TGTCTTCTCAGCATCAAGCAAGG -3' |
| | Reverse | 5'- CCAACACTTCCTGGACAGGCTT -3' |
| ***PERK*** | Forward | 5'- CCAGGCTTTCAAGGTGGACAGT -3' |
| | Reverse | 5'- GGTAGGCAATGAGGACGATGAG -3' |
| | Forward | 5'- GGCTACCATTATCCTGAGCACC-3' |
| ***IRE1 alpha*** | Reverse | 5'- CTCCTTCTGGAACTGTTGGTGC-3' |
| ***XBP1*** | Forward | 5'- TGGACTCTGACACTGTTGCCTC-3' |
| | Reverse | 5'- TAGACCTCTGGGAGTTCCTCCA-3' |
| ***CHOP*** | Forward | 5'- GGAGGTCCTGTCCTCAGATGAA-3' |
| | Reverse | 5'- GCTCCTCTGTCAGCCAAGCTAG-3' |
| ***ATF4*** | Forward | 5'-AACCTCATGGGTTCTCCAGCGA-3' |
| | Reverse | 5'-CTCCAACATCCAATCTGTCCCG-3' |
